# Remodelled Ribosomes Synthesise a Specific Proteome in Proliferating Plant Tissue during Cold

**DOI:** 10.1101/2022.11.28.518201

**Authors:** Federico Martinez-Seidel, Pipob Suwanchaikasem, Dione Gentry-Torfer, Yogeswari Rajarathinam, Alina Ebert, Alexander Erban, Alexandre Augusto Pereira Firmino, Shuai Nie, Michael G. Leeming, Nicholas A. Williamson, Ute Roessner, Joachim Kopka, Berin A. Boughton

## Abstract

Plant acclimation to low temperatures occurs through system-wide mechanisms that include proteome shifts, some of which occur at the level of protein synthesis. All proteins are synthesised by ribosomes. Rather than being monolithic, transcript-to-protein translation machines, ribosomes can be selective and cause effective proteome shifts required for successful temperature acclimation. Here, we use apical root meristems of germinating seedlings of the monocotyledonous plant barley as a model to study changes in protein abundance and synthesis rates during cold acclimation. We measure metabolic and physiological parameters that allow us to compare protein synthesis rates in different physiological states, e.g., in cold acclimation compared to the optimal temperature state. We show that ribosomal proteins are independently synthesised and assembled into ribosomal complexes in root proliferative tissue, and assess how the ribo-proteome shifts during cold may be associated with changes in synthesis and accumulation of macromolecular complexes. We demonstrate that translation initiation is the limiting step during cold acclimation and based on our data propose a model of a ribosomal code that depends on a reconfigured ribosome population, where as a mode of cold acclimation, specific ribosomal protein compositions may confer selective association capabilities between 60S subunits and 48S initiation complexes.

## Introduction

Cold acclimation is a physiological challenge for sessile plant organisms (***Hincha and Zuther, 2020***; ***Middleton et al., 2014***; ***Thomashow, 1999***). Next to extensively studied transcriptional responses (***Seki et al., 2002***; ***Jaglo-Ottosen et al., 1998***; ***Thomashow, 1999***; ***Fowler and Thomashow, 2002***; ***Hincha et al., 2012***), cold-triggered translational reprogramming moves into the focus as a hub for temperature acclimation (***Garcia-Molina et al., 2020***; ***Beine Golovchuk et al., 2018***; ***Schmidt et al., 2013***). Translation provides a second regulatory layer that integrates signals from cellular processes and generates a cold acclimated proteome based on the primary layer of a remodeled transcriptome (***Yu et al., 2020***; ***Martinez-Seidel et al., 2021a***,c; ***Cheong et al., 2021***).

The functional importance of translation is evident in germinating barley seedlings when they stop growing during the first days of acclimation to a cold shift (***Martinez-Seidel et al., 2021c***) and yet continue to accumulate translation-related complexes in the root tip well above the level of plants grown at optimized temperature (***Martinez-Seidel et al., 2021c***). In the absence of growth, synthesis of new ribosomes appears activated or alternatively, the existing ones are less degraded. Regardless of the mechanism, the accumulation of translation-related complexes correlates with evidence of a cold-induced total proteome shift in root proliferative tissue. This shift comprises not only evidence of the accumulation of structural ribosomal proteins (rProteins) but also of elements of ribosome assembly and translation initiation (***Martinez-Seidel et al., 2021c***).

In Arabidopsis, the ribosome population that accumulates after a cold-shift in roots has altered ribosomal protein (rProtein) composition compared to temperature-optimised control conditions (***Cheong et al., 2021***; ***Martinez-Seidel et al., 2021a***), suggesting that temperature-shifts trigger ribosome heterogeneity. Ribosome heterogeneity can contribute to proteome shifts by selective translation of the transcriptome. Targeted translation of mRNAs based on altered ribosomal structures is referred to as ribosome specialization (***Genuth and Barna, 2018***; ***Martinez-Seidel et al., 2020***; ***Slavov et al., 2015***). Functional specialization of ribosomes entails translational events that drive a response to an external (***Lambers, 2022***; ***Berková et al., 2020***; ***Moin et al., 2021***; ***Appels et al., 2021***) or a developmental (***Norris et al., 2021***; ***Shrestha et al., 2022***; ***Xiong et al., 2021***) cue, for example, the preferred translation of specific groups of proteins that facilitate acclimation to low temperatures. These concepts suggest that the previously observed abundance changes of the translational machinery in cold-acclimating root tips of barley seedlings (***Martinez-Seidel et al., 2021c***) may indicate the production of ribosomes tuned to cold-specialized translation. Protein biosynthesis, including ribosome biogenesis, is the largest energy expenditure in the cell (***Ingolia et al., 2019***; ***Russell and Cook, 1995***; ***Verduyn et al., 1991***) and consequently building a specialized ribosome population *de novo* would consume much time and resources. In plants, cytosolic ribosomes have an rProtein-based half-life of 3-4 days (***Salih et al., 2020***), and thus ribosome biogenesis alone may fail to satisfy the demand for customized ribosomal complexes during cold. Thus, there may be a ribosome remodeling mechanism that relies on increased biogenesis and/or *in-situ* ribosome remodelling to alter steady proteome dynamics by causing a cold-induced proteome shift.

The cellular proteome is highly dynamic in sessile plants. The transition between different proteome states is a constant feature in the plant life cycle and varies between organs and tissues (***Nelson and Millar, 2015***). Plant roots contain cells at multiple different developmental and ontological stages that coexist along longitudinal and radial axes (***Dinneny and Benfey, 2008***). The proteome of the root tip is highly dynamic and demonstrably differed from the longitudinal adjacent older tissue, specifically considering the cold triggered abundance changes of translation- and ribosome-associated proteins (***Martinez-Seidel et al., 2021c***). Root apical meristems are “hotspots” of growth and require a high amount of newly synthesised ribosomes to support cell proliferation compared to the adjacent elongation and differentiation zones (***Clowes, 1958***; ***Verbelen et al., 2006***). The relatively large size of barley roots makes them an ideal system to compare proteome dynamics in response to environmental cues. Barley roots allow for feasible sampling of a sufficiently large tip enriched for meristematic cells. Such sampling circumvents one of the major limitations in analysing complex root landscapes as it avoids pooling of highly diverse root zones and the masking of phenomena linked to rapidly proliferating cells by the more abundant static root zones.

The proteome dynamics are related to protein turnover, which is the balance between protein synthesis and degradation. These two processes lead in different contexts to shifts in the relative abundances of active protein pools. Translation is related mainly to protein biosynthesis and approaches to study translational dynamics typically involve the use of stable isotope mass spectrometry as an empirical measure of protein synthesis (K_*S*_) and degradation (K_*d*_) (***Nelson et al., 2014b***,a). The calculation of plant protein K_*S*_ using a stable isotope pulse depends on several parameters that must be considered (***Ishihara et al., 2015, 2021***; ***Li et al., 2017***). The amount of protein fraction that has taken up the externally supplied stable isotope tracer can be determined by isotopolog analysis of proteinogenic amino acids from isolated and hydrolyzed protein fractions or individual purified proteins. Alternatively, proteomics technology allows multiparallel isotopolog distribution analysis of peptides obtained by digestion of complex protein preparations. The time of stable isotope pulse to sample collection allows conversion of observed isotope enrichment and protein concentration values into rates. To prevent physiological differences in plant tissue and environmental factors from confounding these calculations, several variables are required to correct and normalize protein K_*S*_ rates. The first two variables are protein content and relative growth rate (RGR), which are required to adjust K_*S*_ rates between plant systems that differ in these characteristics. A third set of variables describes the dynamics of tracer incorporation into soluble amino acid pools, and label incorporation into these pools may differ between the physiological conditions being compared and between individual proteinogenic amino acids (***Ishihara et al., 2021***). All of these variables are necessary to correct the isotopic envelopes of proteins or digested peptides based on knowledge of their individual amino acid sequences. These corrections are particularly important when the compared plant systems are transitioning between physiological quasi-stable states, as is the case in most studies of development, physiology, and environment x genome interactions, and especially during the well-described sequences of cold acclimation. The non-acclimated and acclimated states of plants drastically alter protein accumulation (***Martinez-Seidel et al., 2021c***), growth dynamics (***Beine Golovchuk et al., 2018***; ***Martinez-Seidel et al., 2021a***,c), and pools of soluble amino acids (***Kaplan et al., 2004***; ***Guy et al., 2008***). Such changes could strongly bias the observed protein K_*S*_ rates without correction.

In this study, we explore the proteome dynamics in barley root tips by stable isotope tracing and proteomic mass spectrometry of a protein fraction enriched for assembled translation-related complexes. We aim to provide new insight into non-steady-state translation dynamics of this rapidly developing plant tissue. We calculate K_*S*_ rates of individual peptides from samples of barley seedlings germinated in the dark prior to emergence of photosynthetic activity and compare acclimation to sub-optimal temperature (4°C) with optimal temperature (20°C). Our experiments correct for differential growth and protein accumulation and validate the calculation of K_*S*_ rates by morphometric and metabolic phenotype analyses. We monitor protein abundances, RGR, and label incorporation into soluble amino acid pools and thereby correct for expected differential isotope tracer dilution of the peptide building blocks. Using the corrected K_*S*_ rates, we investigate whether the ribosome population that accumulates during cold acclimation consists of *de novo* synthesised rProteins or alternatively, whether rProteins remain non-labelled and are re-used. We report changes of rProteins in cold-acclimated ribosomal populations and compare these findings to various other co-purified cellular protein complexes that are preferentially synthesised during cold and are either functionally related to the protein biosynthesis machinery or have different cellular functions.

## Results

### Experimental design

To ensure legitimate, physiological rearing conditions for barley seedlings, we followed previously published procedures (***Martinez-Seidel et al., 2021c***) with few modifications (Figure 1). Seeds were imbibed for 14 hours under sterile conditions, then transferred to liquid medium and germinated on plates for another 48 hours. At this time, we applied different temperature regimes to mimic an agronomically relevant temperature drop after sowing barley in the field. Germination occurred in the dark for an additional 60 hours, which corresponds to an optimal seeding depth equivalent to the typical period of about 5 days before barley seedlings emerge from the soil in the field (***Kirby, 1993***).

**Figure 1.**
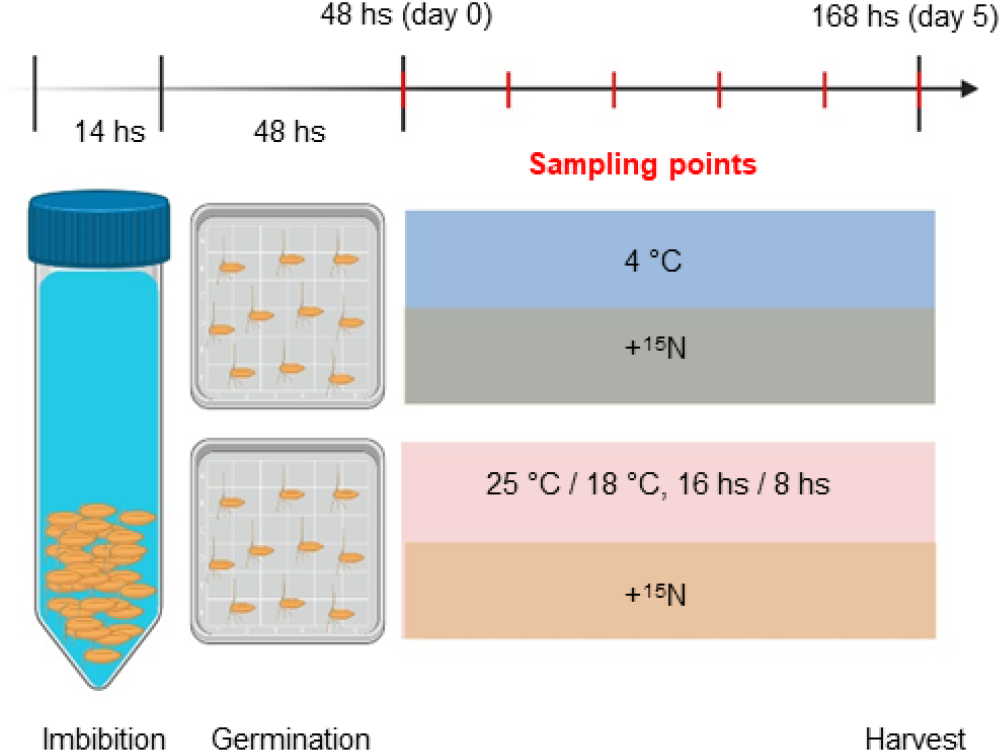
Experimental setup. *Hordeum vulgare* seeds were soaked in sterile H_2_O for 14 h imbibition. Seeds were transferred to plates for germination for 48 h. At 48 h treatments were applied and it was considered as experimental day 0. Half of remaining seedlings were treated with media supplemented with ^15^*N* compounds and the other half continued on media supplemented with ^14^*N* compounds as a control. Half of the seeds in treatment and in control plates were shifted to 4 °C to induce cold acclimation. The other half remained in the control growth chamber with a temperature fluctuation of 25 °C for 16 h and 18 °C for 8 h. Six seedlings were harvested daily per treatment after day 0 for phenotypic analyses. After harvest, each seedling was scanned for phenotyping, roots were weighed for fresh weight and dried for 70 h at 70 °C and weighed again for dry weight. For primary metabolome analysis, root tips were collected in 1.5-cm segments on the fifth day of acclimation. Created with BioRender.com **Figure 1–Figure supplement 1**. Root phenotype during germination assay in optimal control and suboptimal cold temperatures. **Figure 1–Figure supplement 2**. Summary of the methodological workflow to achieve measurements of protein synthesis and abundance in barley root tips.

Control seedlings continued to germinate at an optimal temperature of 25°C/ 18°C in a 16-hour day/8-hour night regime (Figure 1 - Figure S1) with an average temperature of 22°C over 24 hours. Barley seedlings treated with low temperatures were cultured at a suboptimal, but not lethal, temperature of a constant 4°C. Differential growth, one of the necessary variables for calculating individual protein synthesis rates, was monitored by high-resolution imaging of independently germinated seedlings at 12-hour intervals (Figure S1). Throughout the approximately 5-day period, etiolation, premature greening, seed nutrient starvation, and other processes resulting from unnaturally long darkness were avoided (***Kirby, 1993***). Seedlings at 4°C developed more slowly but showed no other macroscopic phenotypic changes (Figure S1). We began labeling seedlings with ^15^N coincident with the temperature shift 48 hours after germination (t_0_) and throughout the differential temperature treatment to ensure that the dynamics of tracer incorporation reflected the physiological change associated with cold acclimation and delayed growth. After 108 hours of germination, i.e. at the last experimental time point (t_*f*_) of the labeling experiments, the root tips of the seedlings were harvested to analyze the isotope incorporation and pool sizes of soluble amino acids and to measure the relative abundances and ^15^N enrichment of individual proteins (Figure 1 - Figure S2).

### Root growth dynamics

Root systems of germinating barley seedlings were carefully phenotyped using non-destructive and paired destructive measurements to characterize differential growth and biomass dynamics induced by different temperature regimes. We expected a strong effect on growth based on previous observations of cold-acclimated *Arabidopsis thaliana* rosette plants delaying growth for a period of 7 days after switching to suboptimal temperatures (***Beine Golovchuk et al., 2018***; ***Martinez-Seidel et al., 2021a***). Such a growth difference can be a confounding variable when comparing protein synthesis rates of acclimated and non-acclimated plants. For example, a non-acclimated plant may produce more protein X (Px) in absolute terms than a cold-acclimated plant, but growth will also differ and is likely to be lower in a cold-acclimated plant. Whether a non-acclimated or a cold-acclimated plant preferentially synthesises Px can only be determined by normalizing biomass accumulation.

We selected an optimized growth measure from several variables that described morphological phenotypes of roots from germinating barley seedlings (Figure 2, Figure 2 - Figure S1 to S2 & Table S1). To this end, we monitored root growth at 12h intervals between t_0_ and t_*f*_ across the germination period.

**Figure 2.**
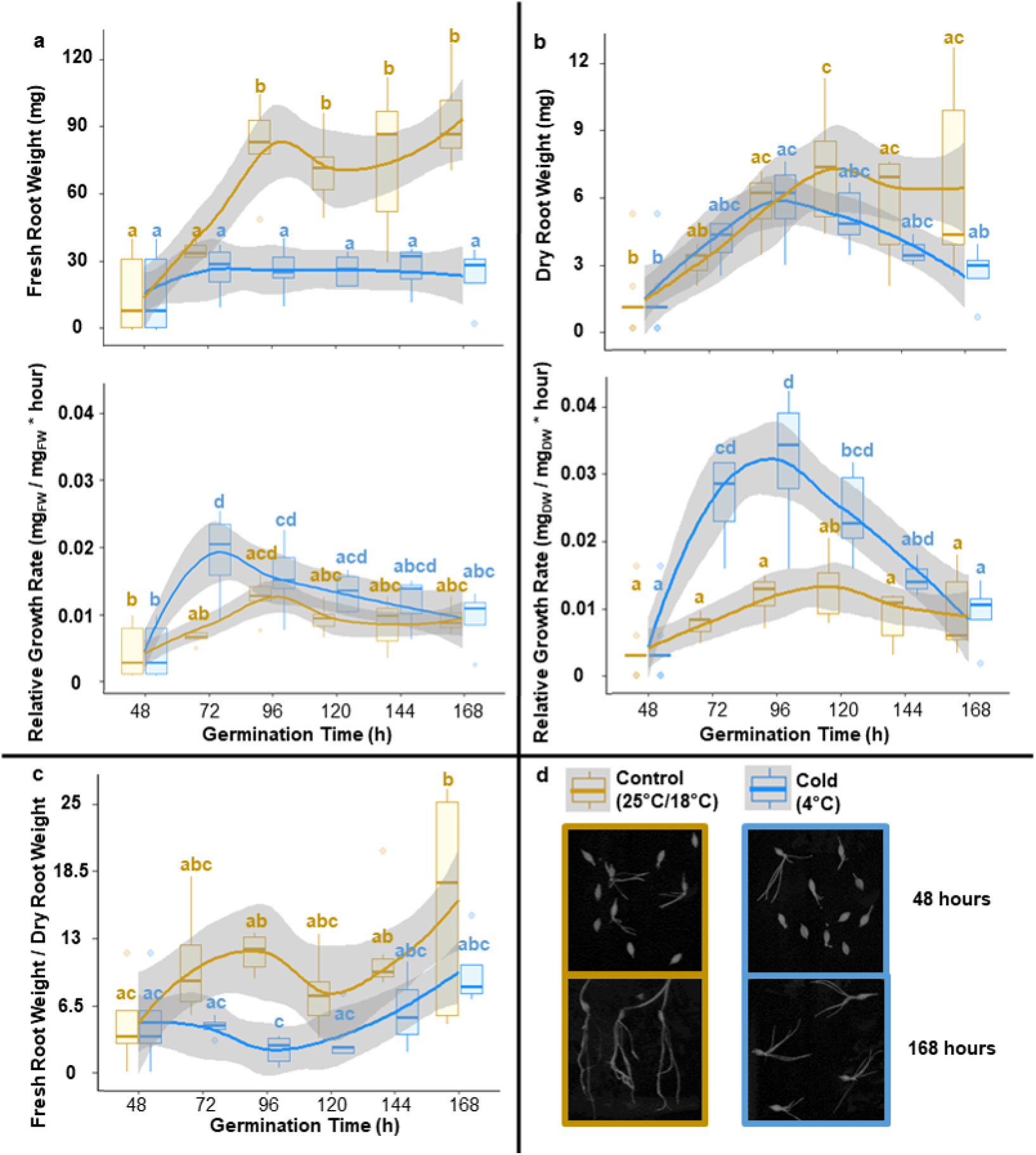
Root growth dynamics of barley seedlings reared at optimal and cold suboptimal temperatures. Barley seeds were imbibed and germinated for 48 hours at optimal rearing conditions and then were transferred to different temperature regimes for five days. Panels A, B, & C contain measurements of specific growth related variables measured in roots, outlined as plots where means are solid coloured lines (blue for cold and gold for control) and standard deviations are shades around the mean. All mean values were compared using an ANOVA followed by the posthoc Tukey HSD test. The boxplots are labelled according to significant differences in the Tukey HSD test, where shared letters indicate lack of significance. Starting at 48 hours after imbibition, seedlings were scanned every 24 hours to measure root growth with a subsequent destructive harvesting to measure root fresh weight (A - upper panel). Subsequently root systems were dried for 3 days at 70°C and weighed again to measure root dry weight (B - upper panel). The recorded weights were used to asses the statistical changes in the fresh to dry weight ratio during the experimental period (C). Finally, both weights were used to calculate RGRs (A & B - lower panels) based on root weight at the imbibition stage (i.e., 0 hours) being zero grams. RGR would subsequently serve the purpose of normalizing protein synthesis rates to basal root growth, preventing biomass accumulation biases. **Figure 2–Figure supplement 1**. Summary of 95% confidence level Tukey-HSD statistical differences in mean levels of growth related variables across treated and control barley seedlings. **Figure 2–Figure supplement 2**. Statistical differences in mean levels (95% confidence level Tukey-HSD) of growth related variables derived from scanning treated (blue) and control (gold) barley seedlings at each time-point. **Figure 2–Figure supplement 3**. Linear regression after natural logarithm transformation of growth related variables.

The roots of the control seedlings accumulated significantly more fresh weight (FW) compared to the roots of the cold-acclimated seedlings within the monitored period between 48h (t_0_) and 108h (t_*f*_) after germination (Figure 2A above). FW accumulation ceased after 60 hours and subsequently remained significantly reduced in the cold-shifted group compared to the control group. In contrast, dry weight (DW) increased equally in the cold-shifted group and the control group, increasing significantly at 72 hours compared with 48 hours (Figure 2B above). After 72 hours, DW was constant. The cold-shifted group appeared to reduce DW, but this observation did not remain significant during the observation period. Analysis of the FW/DW ratio (Figure 2C) showed a significant difference between the cold-acclimated and control roots after 72 hours and an overall reduced ratio in the cold-acclimated roots.

Based on these observations of complex dynamic changes, we determined the RGRs by FW (Figure 2A below) and DW (Figure 2B below) at five intervals (N_*t*_) relative to the average final root mass at t_*f*_ of the cold-shifted and control groups. We used equation 1.

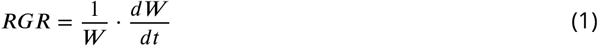

where W is the weight or weight proxy at time (t_*f*_), *dW* is the weight change at time *dt*, which is the time after germination in hours. Consequently, RGR_*F W*_ and RGR_*DW*_ have units mg * mg(t_*f*_)^−1^ * h^−1^, or in short h^−1^. For the RGR calculation at 48h (t_0_), i.e., the start of germination before the temperature shift, we set the root mass at 0h to zero. RGR_*F W*_ and RGR_*DW*_ of the control group were constant during the observation period of 48-108h with small, non-significant fluctuations. While the control group grew steadily, RGR_*F W*_ and RGR_*DW*_ of the cold-transferred group peaked at 60 and 72h, respectively. In this case, we expected transient growth, particularly a decrease in RGRs, due to the transition between optimized and suboptimal temperature regimes. Instead, we found a transient increase in the roots of the cold-acclimated seedlings, which can be explained by the initial DW accumulation after the temperature shift and the partial compensation by the fluctuating water content (Figure 2).

In previous studies on Arabidopsis seedlings, RGR was derived as the slope from log-linear regressions of growth-related variables over time (***Ishihara et al., 2015***). These systems satisfied the assumption of linearity with correlation coefficients (r^2^) approaching 1. For roots of germinating barley seedlings, we had to reject the linearity assumption with r^2^ less than 0.5 for all observed variables, including root length, diameter, volume, length * volume^−1^, number of tips, or branching, as well as FW and DW (Figure 2 - Figure S3). To account for the obviously different RGRs of control and cold-acclimated roots, we chose the average of RGR_*DW*_ over the experimental ^15^N labeling period, 48h (t_0_) - 108h (t_*f*_), for the required normalization (Table 1). Since protein synthesis directly contributes to DW accumulation, RGR_*DW*_ is the most relevant option for correcting protein synthesis rates of two experimental systems that differ in their growth rates. However, DW determination (as well as FW determination) is destructive and requires a significant sample mass. For these reasons, ^15^N-label incorporation analyses cannot be directly paired and require additional replicated experiments. We investigated the potential of non-destructive methods for RGR determination but were unable to find a suitable replacement for RGR_*DW*_ (Table 1). Averaged RGRs by root length and length * volume^−1^ reflected accelerated RGR_*DW*_ in the cold acclimation condition, however, they did not accurately represent the excessive transient increase in RGR_*DW*_, but rather corresponded to what is observed in RGR_*F W*_ (Table 1).

**Table 1.**
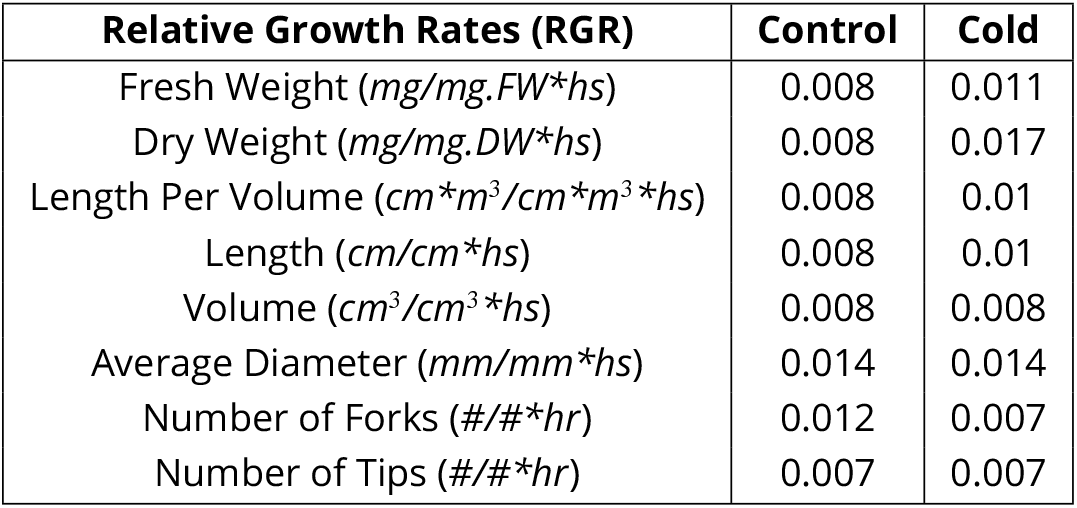
Relative growth rates calculated from multiple root growth proxies from germinating barley seedlings

| **Relative Growth Rates (RGR)** | **Control** | **Cold** |
| --- | --- | --- |
| Fresh Weight (*mg/mg.FW\*hs*) | 0.008 | 0.011 |
| Dry Weight (*mg/mg.DW\*hs*) | 0.008 | 0.017 |
| Length Per Volume (*cm\*m<sup>3</sup>/cm\*m<sup>3</sup>\*hs*) | 0.008 | 0.01 |
| Length (*cm/cm\*hs*) | 0.008 | 0.01 |
| Volume (*cm<sup>3</sup>/cm<sup>3</sup>\*hs*) | 0.008 | 0.008 |
| Average Diameter (*mm/mm\*hs*) | 0.014 | 0.014 |
| Number of Forks (*#/#\*hr*) | 0.012 | 0.007 |
| Number of Tips (*#/#\*hr*) | 0.007 | 0.007 |

### Reprogramming of the Primary Metabolome

Primary metabolism is fundamentally reprogrammed during temperature acclimation (***Kaplan et al., 2004***; ***Guy et al., 2008***). The cold-induced reprogramming involves central metabolism, the source of amino acid building blocks for translation. Therefore, we characterized the primary metabolome of root tips of germinating barley seedlings in the cold-acclimated and non-acclimated condition at t_*f*_ (Figure 3). Cold acclimation contributed most to the overall variance of the resulting multidimensional metabolic data set (Table S2). Principal component analysis (PCA) assigned 65% and 75% of the explained variance to PC1 of the autoscaled or non-scaled data, respectively (Figure 3C-D). The combined technical and biological variance between replicates evident in PC2 was minimal (Figure 3A). In addition to fructans and organic acids, amino acids contributed most to the variance caused by cold acclimation, as inferred from the measure of variable importance in PCA (Figure 3B). Additionally, the externally supplied ^15^N tracer dilutes and distributes differentially among the proteinogenic amino acids, which accumulate in the cold with log_2_-fold changes of approximately 2-50 (Table S2). These observations highlight the need to monitor and correct for ^15^N enrichment in each of the soluble proteinogenic amino acid pools, ideally in split and paired samples of the protein or peptide ^15^N enrichment assays. The amino acids that contributed most to the PCA separation of the primary metabolome from cold and control samples were pyroglutamic acid (derived from glutamine converted via our extraction/derivatization procedures), cysteine, serine, homocysteine, glutamine, glycine, and glutamic acid, all of which were among the top 20 log_2_-fold changed metabolites between conditions.

**Figure 3.**
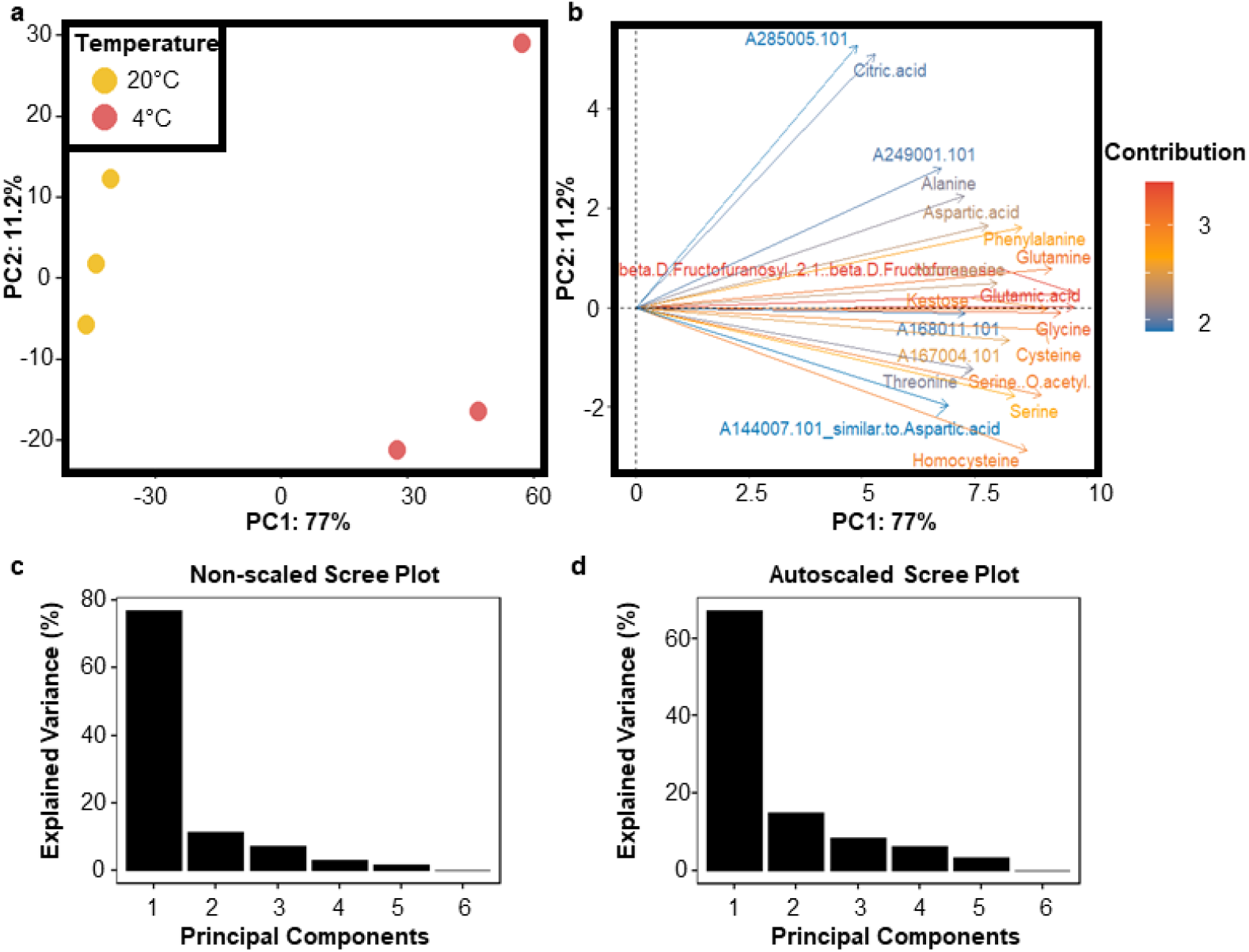
Primary metabolome dynamics of barley seedlings reared at optimal and cold suboptimal temperatures. Also relates to Table S2. The soluble primary metabolome was obtained from frozen and ground plant tissue by methanol / chlorophorm extraction, and the metabolite extracts were chemically derivatized using trimethylsilyl groups to enhance volatility across the gas chromatographic column. Subsequently, the metabolome was measured in a multiplexed array by GC-EI-ToF-MS and GC-APCI-qToF-MS (***Erban et al., 2020***) in technical triplicates. Metabolites were manually annotated in TagFinder and representative tags for each metabolite chosen. Primary metabolome data were analyzed with functions of the RandoDiStats R package. Panel A contains the principal component analyses (PCA) plot where experimental samples are variables and metabolites eigenvectors determining the separation of the samples. Panel B highlights the best 20 metabolite features contributing to the separation outlined in the samples by a contribution PCA plot. Panel C & D contain the scree plot of variance explained in each principal component (PC) in the non-scaled and autoscaled dataset respectively.

### Tracer Dynamics in Soluble Amino Acid Pools

In this study, we used free proteinogenic amino acids as a proxy for estimating the labelling of aminoacyl-tRNAs and monitored the ^15^N-tracer dynamics of the soluble amino acid pools in root tips at t_*f*_ (Figure 4). We supplied a mixture of 99% ^15^N-labelled glycine and serine to obtain a rectangular stable isotope pulse. We chose not to use inorganic ^15^N tracers, which slowly label soluble amino acid pools (***Nelson et al., 2014b***). Inorganic ^15^N labelling arguably leads to low ^15^N incorporation, especially under reserve mobilization conditions of a germinating barley seedling (cf. Discussion). Consistent with our labelling strategy, soluble serine and glycine retained most of the ^15^N tracer within root tips at 15% ^15^N enrichment of the two amino acids in the cold-shifted group and at 2.5 - 5.0% in the control group (Figure 4, Table S3, File S1). The ^15^N enrichment values represent the experimental ^15^N incorporation after correction for the natural isotopic abundances of the elements (NIA). The large difference between conditions clearly indicates that additional correction between experimental conditions is required to allow comparison.

**Figure 4.**
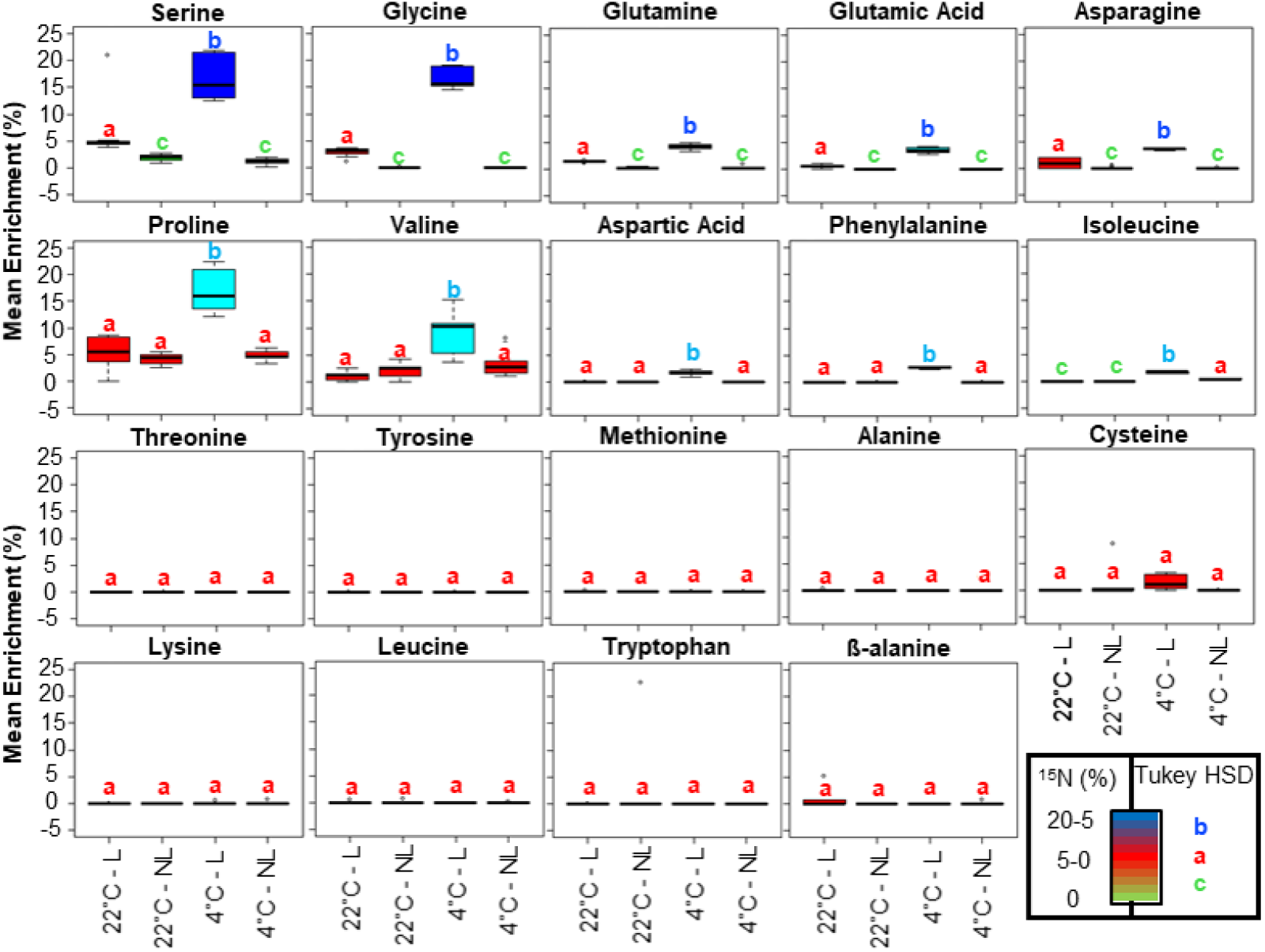
Mean isotopic enrichment of amino acid soluble pools in barley root tips from seedlings germinated at suboptimal (4°C) and optimal (20°C) temperatures. Also relates to Table S3 and File S1. The soluble primary metabolome was obtained exactly as described in Figure 3. Subsequently, at least three independent mass fragments per amino acid analyte along with their isotopolog peak intensities were extracted from total ion chromatograms. The selected fragments for ^15^*N* enrichment percentage calculations were those that appeared in spectra containing less co-eluting ions as well as those with evidently increased mass accuracy (e.g., from the APCI platform) summing up to less noisy spectra. The fragments were corrected for natural isotopic abundance, enabling calculation of enrichment percentages using the R package IsoCorrectoR (***Heinrich et al., 2018***). Finally, mean enrichments were statistically compared across treatments using an ANOVA, followed by a posthoc Tukey HSD test. Boxplots are coloured according to mean significant differences, further emphasised by the letters above each box.

Further correction is required because the ^15^N tracer was incorporated to different extents among the different proteinogenic amino acid pools under the experimental conditions. We monitored 18 proteinogenic amino acids (Figure 4). Histidine and arginine were below the detection limit in our current study. Under control conditions, glutamine, estimated by GC-MS profiling proxy, pyroglutamic acid, glutamate, and asparagine were significantly labelled at 2.5 to 5.0% ^15^N enrichment. The cold-shifted root tips picked up to 2.5-15.0% ^15^N labelling in glutamine, glutamate, asparagine, and additionally in proline, valine, aspartate, phenylalanine, and isoleucine (Figure 4, Table S3). The remaining monitored proteinogenic amino acids and beta-alanine, a non-proteinogenic amino acid control, did not absorb ^15^N. Our analysis provided accurate ^15^N enrichment of most proteinogenic amino acids that could be paired with protein and peptide enrichment measurements. Because even minor contributions from multiply labelled amino acids add up to substantial ^15^N incorporation into peptides, failure to account for these contributions may introduce bias into estimated protein synthesis rates. This confirms the need to study the dynamics of the internalized tracer.

### Protein Synthesis during Transition from a Physiological Steady State

Throughout their lifespan, plants constantly transition through physiological and proteomic states that adapt to developmental and environmental factors (***Nelson and Millar, 2015***). Finding conditions that approach a steady state and allow assessment of protein turnover using stable isotope mass spectrometry to track both synthesis rates (K_*S*_) and degradation rates (K_*d*_) of specific proteins is no easy task. Like many other plant systems that respond to stressors, the root of the barley seedling acclimated to cold is constantly changing (Figure 2) and did not reach an equilibrium state within our observation period. Therefore, we determined K_*S*_ over the observed time interval rather than turnover.

We based our calculations on two published strategies used to determine K_*S*_ from turnover calculations of plant proteins (***Ishihara et al., 2015***; ***Li et al., 2017***). Both of these studies normalize K_*S*_ of individual proteins using growth and protein accumulation rates, as described in the Methods section by equations 3a & 3b (***Ishihara et al., 2015***) and 4a, 4b, 4c & 4d (***Li et al., 2017***). Specifically, we determine the ^15^N enrichment of peptides, consider ^15^N enrichment in soluble proteinogenic amino acid pools, and correct K_*S*_ by relative growth transformed into relative protein accumulation rates. Equation 2a calculates K_*S*_ as the product of labelled peptide fraction (LPF) times a modified version of RGR (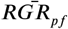 in equation 2b) times a factor of 100, which ensures that K_*S*_ units are expressed as a percentage (%) of the normalized labelled peptide fraction accumulated per unit of protein weight per hour. (equation 5a).

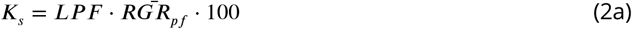

Since our study is not at a biological steady state, we introduced the modified 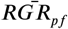 calculation. First, we calculate the average RGR_*DW*_ over the labelling period 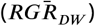, i.e., the observed sum of the measured RGR_*DW*_ of the analyzed time intervals between t_0_ and t_*f*_ divided by the number of time intervals (N_*t*_), replacing *dW* (weight accumulation at time t) with P_*f*_ (equation 2b).

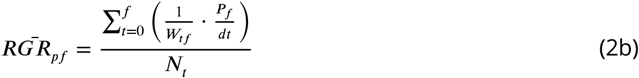

The factor P_*f*_ is used to convert the RGR_*DW*_ into a relative accumulation rate of total protein with respect to DW (RGR_*pf*_). To this end, we determined P_*f*_, which is the ratio of the final total protein mass (mg [(P_*pf*_]) to the dry mass at time t (mg [*dW*]), equation 2c. For both empirical measurements in milligrams, this process cancels the units and converts them to a fraction of the total final protein relative to the cumulative weight.

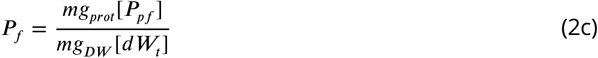

To calculate the labelled peptide fraction (LPF, equation 2d), we extracted the distributions of the mass isotopologs from the parent ion masses of LC-MS/MS data sets. We calculated the expected parent ion monoisotopic masses of the peptides based on their molecular formula derived from the known amino acid composition. We determined the isotopolog abundances at mass intervals corresponding to the mass-to-charge ratio of the peptide. These initial isotopolog distributions were corrected for the natural isotopic abundances (NIA) of the elements according to their molecular formula. The resulting NIA-corrected isotopolog distributions exclusively reflected the experimental ^15^N labelling and allowed the calculation of the fractional ^15^N enrichment of peptides, i.e., the ratio of ^15^N atoms in a peptide to the sum of all N atoms. Finally, the fractional enrichment or non-corrected LPF is corrected by a constant that differs for each peptide as a quotient, equation 2d.

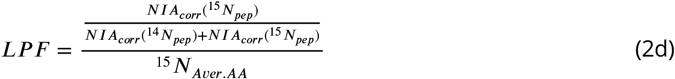

The ^15^*N*_*Aver*.*AA*_ needs to be the quotient dividing non-corrected LPF in order to increase the fractional enrichment in individual peptides, which assumes a fully labelled source, by what is actually the achieved labelling, which is always only fractional given the partial labelling in soluble amino acid pools. Thus, the varying condition-dependent ^15^N incorporation into soluble proteinogenic amino acids and the amino acid composition of peptides determine the maximum fractional enrichment that peptides can achieve. To account for these factors, we correct the fractional ^15^N enrichment of peptides by dividing by a peptide- and condition-specific correction factor (^15^*N*_*Aver*.*AA*_ in equation 2e) to obtain a corrected version of the LPF. ^15^*N*_*Aver*.*AA*_ is the average NIA-corrected fractional ^15^N enrichment over all amino acids in a peptide assuming that the fractional ^15^N enrichment of each amino acid incorporated into protein is equal to its observed fractional ^15^N enrichment in the soluble amino acid pools, Equation 2e. In other words, ^15^*N*_*Aver*.*AA*_ corresponds to the maximum fractional ^15^N enrichment that each peptide can achieve under specific labelling conditions. ^15^*N*_*Aver*.*AA*_ only considers the labelled amino acid residues because these are entered as potentially labelled N atoms to define the combinatorial matrix used to correct for NIA. Thus, unlabelled soluble amino acids have no effect on our calculations.

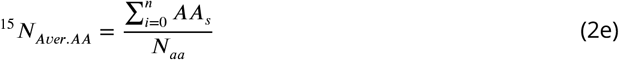

Consistent with an expected immediate rectangular ^15^N label incorporation into soluble amino acid pools, we used the NIA-corrected ^15^N incorporation into amino acids as determined at t_*f*_ (***Ishihara et al., 2015***). We tested an alternative scenario for delayed incorporation by modelling incorporation as a linear function between t_0_ and t_*f*_ (Table S4F and File S2.1 & S2.2). This leads to a more conservative trend of enrichment incorporation. Otherwise, the assumption of rectangular incorporation would mean that the average labelling in soluble amino acid pools over the experimental period is overestimated. Nevertheless, because of the large increase in peptides exceeding the maximum possible LPF (> 1.0), we rejected the alternative assumption of substantially delayed incorporation of label into soluble amino acid pools. Instead, we used the maximum possible LPF as a reference to accept or reject low-enriched soluble amino acids as part of the correction factor and thus be able to decide on the correct amino acid combination.The complete workflow for data mining and computation of our results (Figure 5) is provided by the ProtSynthesis R package, which contains detailed code annotations and descriptions of the computational procedures used in this and previous studies (***Ishihara et al., 2015***; ***Li et al., 2017***). The complete workflow (Figure 5) and its usage details are deposited in two GitHub repositories. Namely, the repository ProtSynthesis, from which the R package can be installed directly in any R environment, and the repository isotopeEnrichment, which contains the usage instructions and the Python function that allows direct mining of peptide isotopolog abundances.

**Figure 5.**
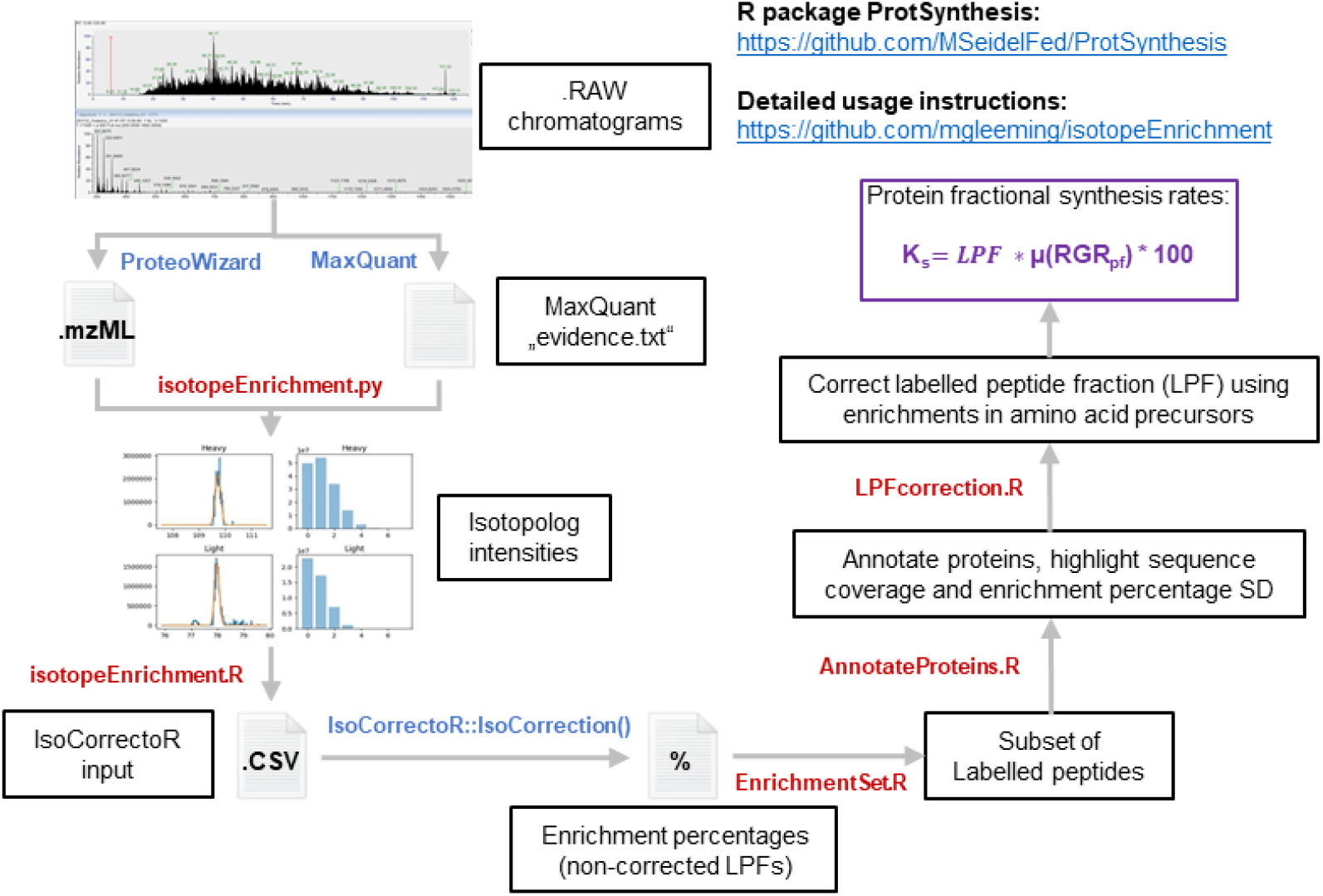
Bioinformatics pipeline written as a python function and an R package that jointly enable to calculate stable isotope tracer incorporation into peptides from LC-MS/MS data and derive fractional protein synthesis rates in the organismal physiological context. Also relates to Table S4. The whole pipeline has been made publicly available in two repositories. The isotopeEnrichment GitHub repository, which contains detailed usage instruction of both python and R functions and the ProtSynthesis R package, which can be installed into any R environment via devtools. All the self-written steps in the pipeline are signalled by red font, while existing algorithms that are external dependencies are depicted in blue font. The pipeline entails using the MaxQuant software to locate proteins and peptides in the chromatograms and then trace back their position and recover their full isotopolog intensities from .mzML files using “isotopeEnrichment.py”. Subsequently, an optimal number of isotopolog peaks are derived for individual peptides based on the molecular formula and the enrichment percentages in soluble amino acids using “isotopeEnrichment.R”. The data delivered is organised as is required by the IsoCorrectoR R package, which is then used to correct for the natural isotopic abundance and calculate the enrichment percentage of individual peptides (LPF). Subsequently, statistical filters are used to identify and annotate significantly labelled proteins as well as to derive relevant statistics that detail the quality of the protein hit using “EnrichmentSet.R” and “AnnotateProteins.R” respectively. Finally, “LPFcorrection.R” is used to correct the calculated LPFs using the enrichment in soluble amino acid pools, with these corrected values fractional protein synthesis rates are calculated by multiplying them by relative growth rates times 100. **Figure 5–Figure supplement 1**. Summary of 95% confidence level Tukey-HSD statistical differences in mean levels of protein content from proteome fractions enriched in ribosomes across treated and control barley seedlings. **Figure 5–Figure supplement 2**. Subset of peptides considered as having optimal quality for interpretation of their relative fractional synthesis rates during the physiological transition of roots from germinating barley seedlings from optimal to suboptimal temperature.

### Synthesis and Accumulation of Macromolecular Complexes during Cold

To validate our method, we obtained a ribosome-enriched fraction of the barley root tip proteome by filtering a cell lysate through a 60% sucrose cushion, recovering the pelleted fraction. This ensured that only assembled macromolecular complexes were monitored. The comparative fractions obtained were from treated and control seedlings. Using the same extraction procedure, the total protein content of cell complexes was significantly higher in the roots of treated plants than in the roots of control plants (Figure 5 - Figure S1 and Table S5). Using the LFQ-normalized protein abundances (***Zhang et al., 2012***) of peptides extracted from the recovered fraction, we then report what fraction of the monitored complexes accumulate in barley root tips at sub-optimal low compared with optimized rearing temperatures (Table 2). At the same time, we monitored the incorporation of ^15^N into the individual protein components of these complexes and found 1379 good quality peptides after applying our method (Figure 5 - Figure S2 and Table S4G), from which we can confidently report fractional synthesis rates. From this information, we can estimate what part of the complexome proteome accumulation is due to protein synthesis and what part is due to the lack of protein degradation (Table 2).

**Table 2.**
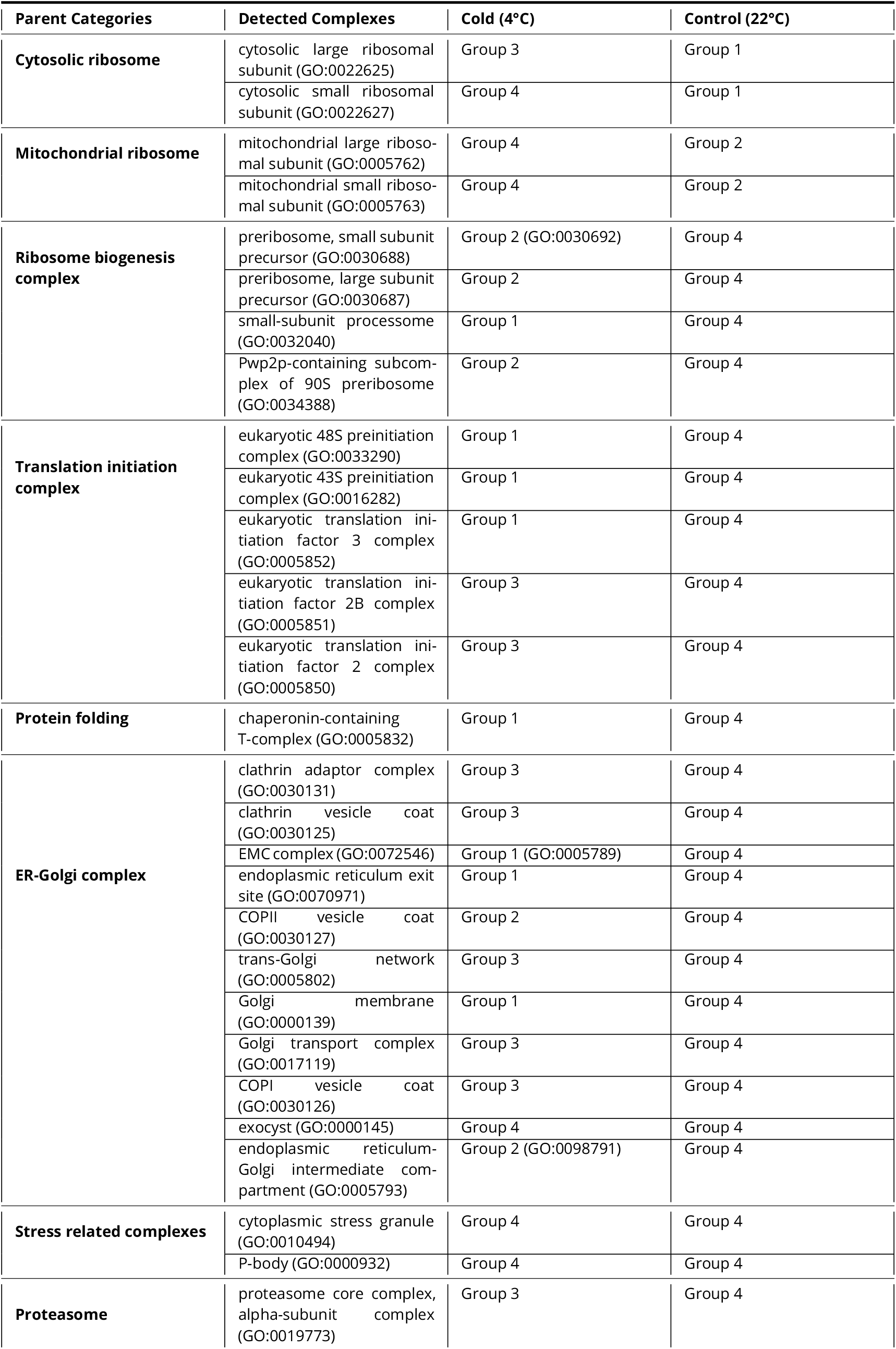
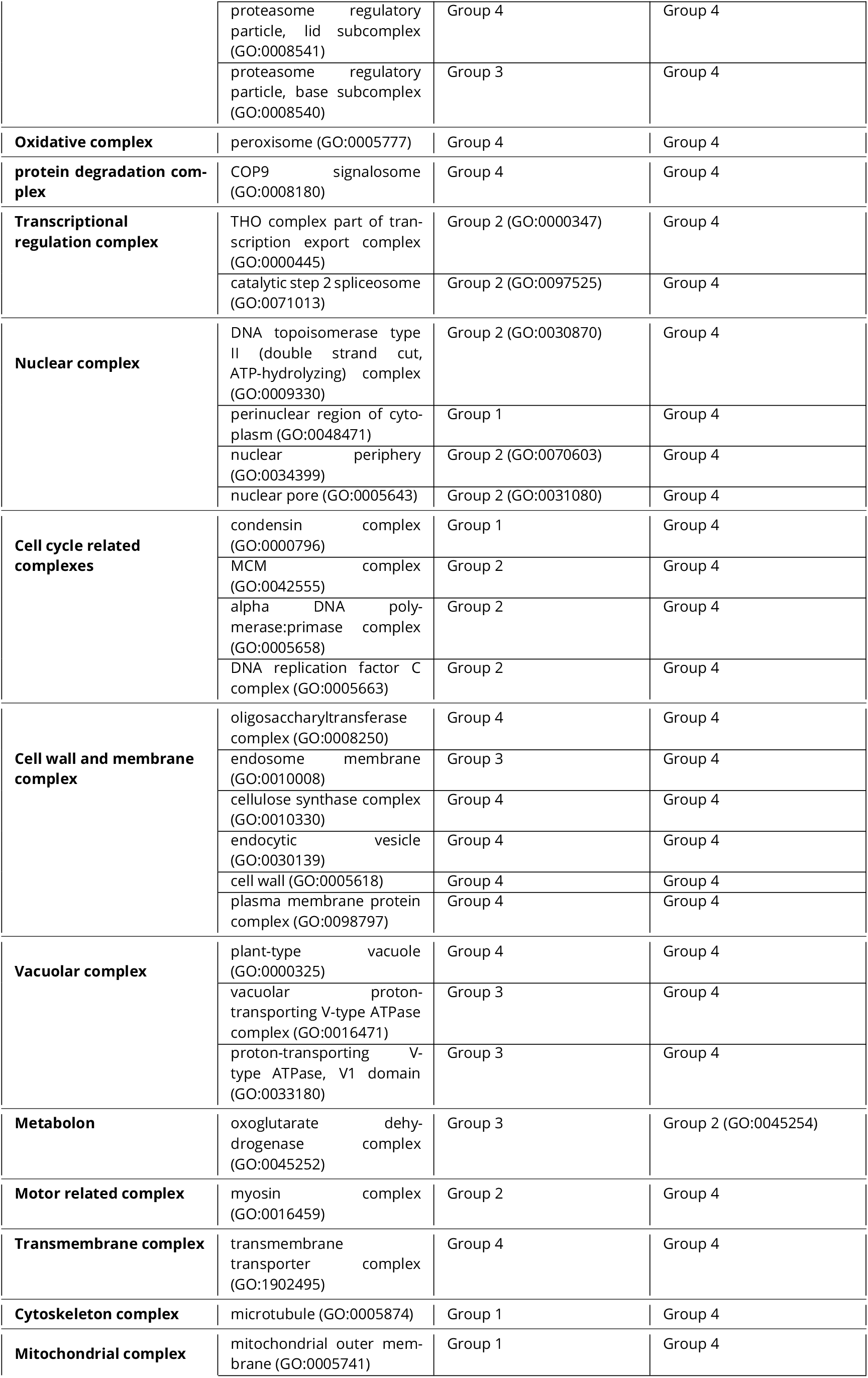
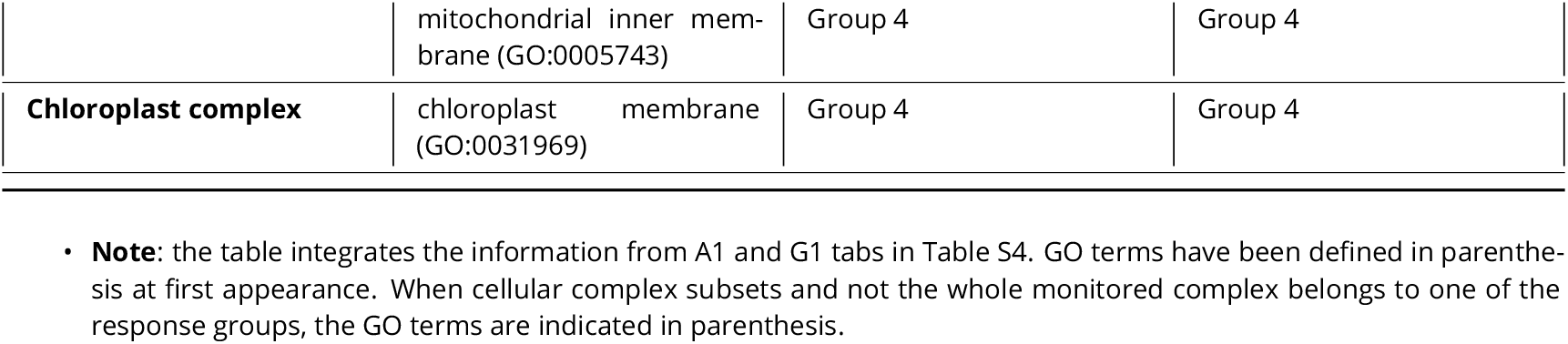
Accumulation and origin of protein components from detected multi-protein complexes in barley root tips classified in four groups of responses during the experimental period: (Group 1) accumulated and newly synthesised, (Group 2) accumulated and not degraded, (Group 3) not accumulated but newly synthesised, (Group 4) and not accumulated and not synthesised.

There are four groups of responses for protein-complexes and their components in Table 2. Group one contains proteins that accumulate and are preferentially synthesised at a specific temperature. Group two contains proteins that accumulate due to lack of protein degradation because they do not incorporate the nitrogen isotope. Group three contains proteins that do not accumulate and nevertheless are preferentially synthesised at a specific temperature, implying high turnover. Group four contains proteins that are detected but do not accumulate nor are preferentially synthesised. The three significant groups (1-3) can be found at sub-optimal low temperature whereas only group one and two are found at optimal temperature.

#### Altered Ribozyme-Mediated Proteome Remodelling

The global ontology term of cytosolic translation in Table 2, which includes ribosome biogenesis, has proteins that belong to the three significant groups of responses (1-3) at both temperature regimes.

The accumulation dynamics of ribosome biogenesis complexes, which give rise to mature and translationally competent ribosomes, provides insight into the origin of assembled ribosomes. For example, the 90S pre-ribosome in the nucleolus leads to pre-60S and pre-40S complexes. Following their maturation, pre-40S complexes are shaped by the small-subunit processome. Protein components from all these four types of biogenesis complexes significantly accumulate at sub-optimal low temperature. Interestingly, the small-subunit processome features proteins from group one, i.e., accumulating due to *de novo* synthesis. Whereas the other three complexes accumulate due to lack of degradation (group two). This implies that the only remodelled complex from the biogenesis subset is the small-subunit processome, while the others keep functioning in the same state as that from plants growing at optimised temperature.

In terms of assembled cytosolic ribosomes, their structural protein components, as well as those from mitochondrial ribosomes, are accumulated preferentially at optimised temperature. The cytosolic ribosome components are preferentially synthesised, which categorises them in group one. In contrast, the mitochondrial ribosome components are not preferentially synthesised and thus accumulate due to lack of degradation (group two). On the other hand, the protein structural components of the cytosolic large ribosomal subunit are preferentially synthesised but not accumulated during cold, which categorises them in group three and indicates a remodelling aspect that is coupled to high turnover of these complexes at sub-optimal temperature.

Translation initiation complexes preferentially accumulate only at sub-optimal low temperature and, all of them belong to group one, where their proteins accumulate due to *de novo* synthesis.

Beyond the translation ontology term, many of the detected protein complexes that preferentially accumulated at low sub-optimal temperature belonged to group one, where their protein components are also newly synthesised. Two major ontological groups stand out as being newly synthesised by ribosomes and accumulated during cold. In brief, cellular machinery to cope with protein misfolding and aggregation is newly synthesised and accumulated (CCT complex protein components and heat shock proteins) as well as cellular complexes that mediate remodelling of the cell walls and cellular membranes. Thus, this prompted us to analyse in more depth the extent of triggered ribosome heterogeneity and its potential structural link to the observed proteome shift that happens due to protein synthesis by ribosomes.

### Recycled and Remodelled Ribosomes Accumulate during Cold Acclimation

To characterise cold-induced rProtein heterogeneity and decipher its origin, we adjusted the protease digestion to the requirements of a highly basic ribosomal proteome (Figure 6 - Figure S1 and Table S6), and used the optimized method to profile ribosome-enriched barley proteome extracts. To validate that our method enabled recovery of native ribosomes, we subjected *Escherichia coli* 70S ribosomes to the same purification method and assessed the completeness of the recovered ribo-proteome. We found 21/21 30S small subunit rProteins and 33/33 50S large subunit rProteins, which featured reproducible abundances in triplicated measurements (Figure 6 - Figure S2 and Table S7). We then calculated protein abundances, relative stoichiometry, and fractional synthesis rates for ribosome structural protein components of the barley extracts at the onset of the physiological transition from optimized germination conditions to sub-optimal low temperatures (Figure 6 and Table S4H).

**Figure 6.**
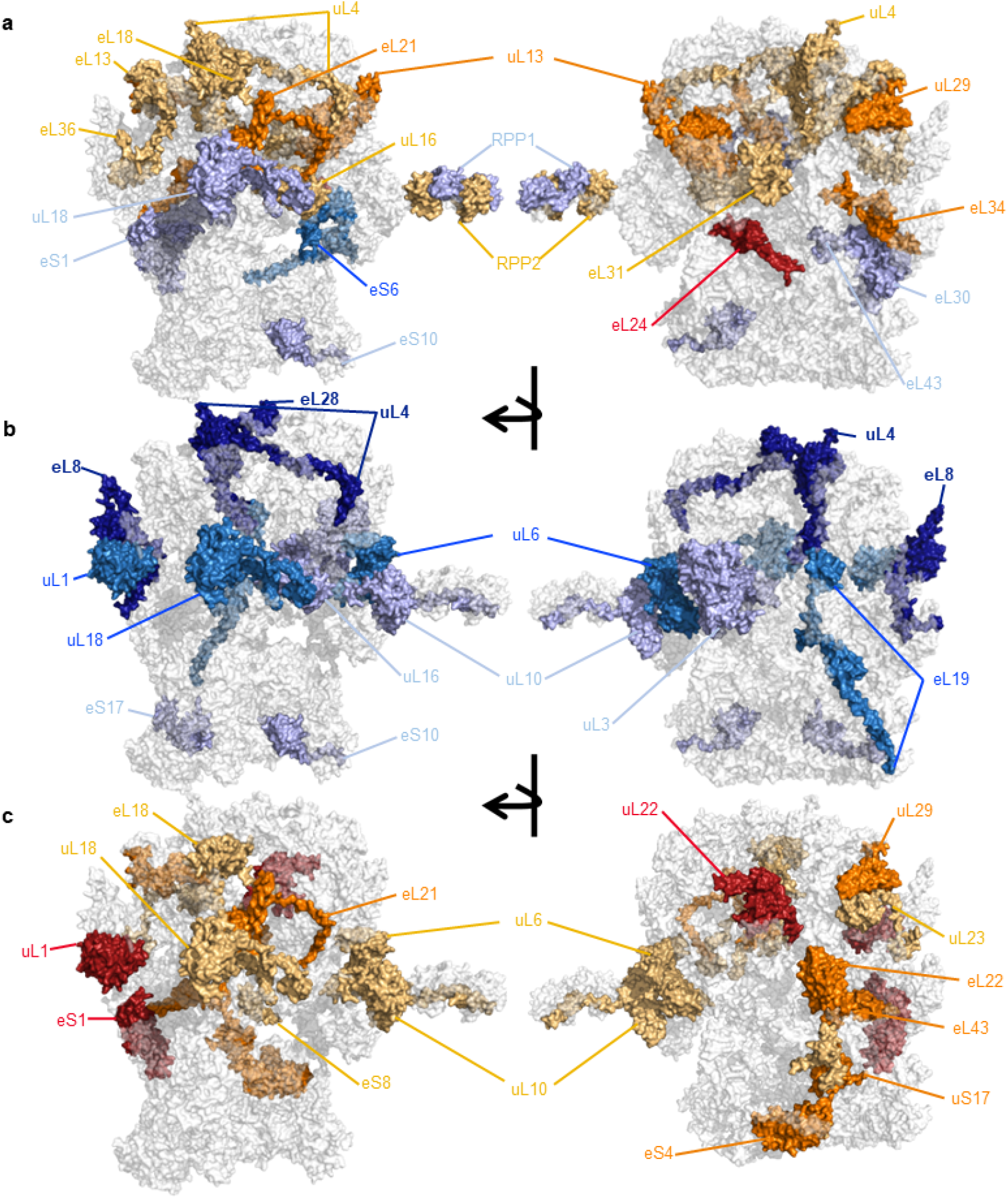
Characterization of cold heterogeneous barley ribosomes, their ribosomal protein (rProtein) composition, fractional synthesis rates and induced substoichiometry. Barley ribosomes from plant ^15^*N* labelled proliferative tissue were purified and used to profile the rProteome. Abundances and isotopolog envelopes were recovered from the Mass Spectrometry data and used to calculate average ribosome relative stoichiometry of rProteins and their fractional synthesis rates. (a) rProtein substoichiometry; orange and reds are rProteins accumulated at optimal temperature and blues rProteins accumulated during cold in the ribosomal population. (b) Preferential synthesis of assembled rProteins during cold acclimation. (c) Preferential synthesis of assembled rProteins during control temperature. Preferential synthesis refers to peptides with significant changes (*P*_*adj*_ values < 0.05, n = 3, tested using a customized generalized linear model) in their fractional synthesis rates. **Figure 6–Figure supplement 1**. Volcano plot outlining the differences in mean peptide length produced with Lys-C or Trypsin, during an *in silico* protease digestion test, as proteases to digest the Arabidopsis signature plant ribosomal proteome. **Figure 6–Figure supplement 2**. Positive control and complete coverage of *escherichia coli* 70S ribosomal proteome from a commercially available preparation used to verify the ribosomal proteomics pipeline. **Figure 6–Figure supplement 3**. Ratio between 40S SSU and 60S LSU abundances across experimental samples

Previously, we observed that rProteins, when averaging ribosome-bound and ribosome-free forms, increase in abundance in barley root tips of germinating seedlings subjected to cold acclimation (***Martinez-Seidel et al., 2021c***). Here, we found reliable LFQ intensities for 17 rProtein families from the small 40S subunit (SSU) and 38 rProtein families from the large 60S subunit (Table S4C), all of which were bound to ribosomal complexes. Among these 55/80 high confidence rProtein families, a total of 95 paralog genes were identified.

#### Ribosomal Protein Substoichiometry

The sum of SSU-rProtein abundances correlated linearly over an r^2^ of 0.98 with the sum of LSU-rProtein abundances across all samples, maintaining a constant ratio of 3x LSU to 1X SSU (Figure 6 - Figure S3 and Table S4C). Nevertheless, the LFQ abundances of all identified paralogs decreased in their ribosome-bound form during cold acclimation, either significantly (Table S4B) or not. Thus, the control seedlings had in average more 40S and 60S assembled subunits in their root tips. Consequently, we used the sum of 40S proteins to normalize the individual rProtein abundances of SSU and the sum of 60S proteins to normalize the individual rProtein abundances of LSU in order to correct for the relative amount of complexes across samples, as was previously reported (***Martinez-Seidel et al., 2021a***). The normalization allowed us to determine whether these complexes exhibit induced substoichiometry in their rProtein compositions (Figure 6A).

The population of 40S subunits was enriched in four rProtein paralogs during cold, namely eS10 (HORVU3Hr1G111760), eS1 (HORVU1Hr1G032060 and HORVU4Hr1G070370), and eS6 (HORVU2Hr1G010870). Thus, the population of 40S subunits in barley root tips is not canonically complete and sometimes these paralogs are absent compared with the cold population. The population of 60S subunits contains paralogs that are relatively depleted or accumulated in cold conditions. The rProtein families P2 (HORVU1Hr1G073640), uL16 (HORVU1Hr1G024710), eL13 (HORVU7Hr1G067060), eL18 (HORVU2Hr1G018820), eL31 (HORVU7Hr1G050170), eL36 (HORVU5Hr1G009600), uL4 (HORVU4Hr1G075710), eL34 (HORVU7Hr1G071240), uL29 (HORVU2Hr1G068120), eL21 (HORVU1Hr1G038890), uL13 (HORVU5Hr1G096060 and HORVU2Hr1G063900), eL24 (HORVU3Hr1G080130 and HORVU5Hr1G111510) are depleted during cold. The rProtein families P1/P2/P3 (HORVU7Hr1G075360), eL30 (HORVU0Hr1G023290), eL43 (HORVU1Hr1G085170), uL18 (HORVU2Hr1G073320) are accumulated during cold.

#### Altered Ribosomal Protein Synthesis and Ribosome Remodelling

We next examined rProtein synthesis rates to understand what constituted the newly synthesised rProtein substoichiometry (Figure B-C). In general, the cold-induced changes in rProtein synthesis did not coincide with substoichiometry, suggesting independence between rProtein synthesis and ribosome assembly/remodeling. The population of 40S subunits was assembled or remodeled using four rProtein paralogs that were significantly more synthesised at optimized temperature, namely eS8 (HORVU6Hr1G056610), uS17 (HORVU2Hr1G067370), eS4 (HORVU1Hr1G021720), eS1 (HORVU1Hr1G032060). Similarly, two rProtein paralogs were significantly more synthesised during cold: eS10 (HORVU3Hr1G111760), eS17 (HORVU6Hr1G056610). The population of 60S subunits was assembled or remodeled using several rProtein paralogs, which were significantly more synthesised at optimized temperature or during cold. Paralogs uL10 (HORVU7Hr1G073720), eL18 (HORVU2Hr1G018820), uL23 (HORVU2Hr1G086360), uL18 (HORVU2Hr1G073320), eL6 (HORVU7Hr1G00206), eL21 (HORVU1Hr1G038890), eL22 (HORVU2Hr1G019160), uL29 (HORVU2Hr1G068120), eL43 (HORVU1Hr1G085170), uL22 (HORVU5Hr1G052280), uL1 (HORVU7Hr1G059090) were preferentially synthesised and assembled at optimized temperature. Paralogs uL16 (HORVU1Hr1G024710), uL3 (HORVU4Hr1G019980), eL19 (HORVU2Hr1G018700), uL1 (HORVU3Hr1G084310 and HORVU1Hr1G085730), uL18 (HORVU5Hr1G092630 and HORVU2Hr1G073320), eL6 (HORVU7Hr1G00206), eL28 (HORVU1Hr1G079370), uL4 (HORVU4Hr1G075710), eL8 (HORVU7Hr1G054670) were preferentially synthesised and assembled in the cold.

From our results, five types of paralog-specific and rProtein family ribosome association and synthesis dynamics can be deduced:

1. **Paralog switches**

- based on protein synthesis, the uL1 paralogs HORVU3Hr1G084310 and HORVU1Hr1G085730 (cold) and HORVU7Hr1G059090 (control).
- based on substoichiometry, P1/P2/P3 paralogs HORVU7Hr1G075360 (cold) and HORVU1Hr1G073640 (control).
2. **Families with paralogs sharing ribosome-bound accumulation or synthesis dynamics:**

- based on protein synthesis, uL18 HORVU5Hr1G092630 and HORVU2Hr1G073320 (cold-specific); uL1 HORVU3Hr1G084310 and HORVU1Hr1G085730 (cold-specific).
- based on substoichiometry, eL24 HORVU3Hr1G080130 and HORVU5Hr1G111510 (control-specific); uL13 HORVU5Hr1G096060 and HORVU2Hr1G063900 (control-specific); eS1 HORVU1Hr1G032060 and HORVU4Hr1G070370 (cold-specific).
3. **Paralog splice variants (peptides from the same protein with different synthesis dynamics under cold and control conditions):**

- uL10 HORVU7Hr1G073720 (one peptide from exon 5 is cold synthesized and one from exon 4 cold synthesized).
4. **Paralogs that share ribosome-bound accumulation and synthesis dynamics:**

- eS10 HORVU3Hr1G111760; eL18 HORVU2Hr1G018820; eL21 HORVU1Hr1G038890; uL29 HORVU2Hr1G068120; uL18 HORVU2Hr1G073320
5. **Paralogs with inverse ribosome-bound accumulation and synthesis dynamics:**

- accumulated in control ribosomes but preferentially synthesised in cold; uL16 HORVU1Hr1G024710; uL4 HORVU4Hr1G075710
- accumulated in cold ribosomes but preferentially synthesised during control; eS1 HORVU1Hr1G032060; eL43 HORVU1Hr1G085170

## Discussion

### Morphological Phenotype of Barley Roots during Low Temperature Germination

At the physiological level, plants exhibit a growth arrest during the first week of cold acclimation (***Beine Golovchuk et al., 2018***; ***Martinez-Seidel et al., 2021a***,c). *Arabidopsis thaliana* roots reduce mitotic division but not cell elongation at a sub-optimal temperature of 4°C, which significantly reduces meristem size (***Ashraf and Rahman, 2019***). Similarly, barley roots from acclimated seedlings reduce their protein content (***Martinez-Seidel et al., 2021c***), which is mainly related to dry weight accumulation and thus by proxy to mitotic division, suggesting that the cold phenotype of barley and Arabidopsis may share this aspect. In our system, the root length and volume of non-acclimated seedlings increased significantly and steadily compared to the acclimated counterparts, which was enhanced by water accumulation as shown by differences in fresh weight. Conversely, root dry weight did not differ between acclimated and non-acclimated seedlings. The ratio of fresh weight to dry weight remained constant at the end of the acclimation period, and yet there was a transient relative increase in dry weight accumulation from the second to the third day causing a significant imbalance in the dry to fresh weight ratio of acclimated seedlings. In consequence, the dynamics of dry weight accumulation during germination of acclimated barley seedlings are not comparable to their non-acclimated pairs and, therefore, global analyses of protein turnover need to account for this difference.

### Cold Metabolic Phenotype and Testable Links to Translational Responses

In spite of not growing, roots from acclimated barley seedlings show a transition to cold-induced metabolic states. After two days of cold treatment (4 °C) in the dark, glucose increases in barley seedlings, while proline, sucrose, and total lipids decrease (***Zúñiga et al., 1990***). Thus, there is a transient decrease in the abundance of these compounds during the early stages of acclimation, coinciding with the peak of dry weight accumulation reported in our study. The transient decrease in metabolites could subsequently attenuate and then increase again at later stages of acclimation. Transcriptional studies in barley leaves on the third day of cold acclimation (3°C during the day, 2°C at night) predict the accumulation of sugars and polyols such as maltose, glucose, trehalose, and galactinol (***Janská et al., 2011***), and we show that glucose and sucrose were accumulated on the fifth day of acclimation (maltose was not accumulated, and galactinol and trehalose were not detectable). In contrast, the amino acid biosynthetic pathway is not predicted to be upregulated at the transcriptional level, and yet we observed that 25 of the 29 amino acids we detected (including the well-known osmoprotectant proline) were accumulated under cold conditions compared with the optimal temperature. Thus, a cold-metabolic state in our system is consistent with evidence for potential translational control that influences the acclimation response by differentially impacting the amino acid biosynthesis pathway. Amino acid pools can increase from mobilised nitrogen resources or from enzymatic synthesis, the latter of which requires the accumulation of specific proteins through direct or indirect translational control, because amino acid biosynthesis is not predicted to be upregulated based on barley cold-transcriptional dynamics (***Janská et al., 2011***). Our report demonstrates that some of the accumulated amino acids also incorporated ^15^N during cold, suggesting that they may be in part newly synthesized and not just degradation products of storage proteins, as will be discussed in the following paragraph. The role of all accumulated soluble sugars, amino acids, and polyols in the context of cold acclimation is thought to be that of an osmoprotector, i.e., compounds that stabilise proteins and membranes and thus contribute to freezing tolerance (***Rontein et al., 2002***).

### Amino Acid Metabolism and ^15^N Isotopic Flux

How nitrogen uptake and supply occurs in germinating barley seedlings determines the best strategy for isotope flux studies in this system. Most of the nitrogen resources used by germinating barley embryos come from degraded storage proteins located in the endosperm during germination (***Ma et al., 2017***; ***Rosental et al., 2014***; ***Nonogaki, 2008***; ***Lea and Joy, 1983***). Thus, nitrogen transport into the embryo in the form of amino acids and peptides is critical for controlling and setting efficient germination. For example, nitrogen transport and reassimilation is fundamental for the development of gene expression programs in barley caryopses (***Mangelsen et al., 2010***). Consequently, germinating barley embryos activate genes involved in biosynthesis, metabolism, and transport of amino acids at an early stage of 2 to 3 days after germination (***Sreenivasulu et al., 2008***), with peptide transporters considered particularly critical for normal germination processes (***Waterworth et al., 2000***). At the proteome level, nitrogen mobilization systems that supply nutrients to the growing embryo are induced and activated (***Osama et al., 2021***). For example, proteases, including carboxypeptidases and aminopeptidases, provide peptide or amino acid substrates that are released, transported, and used during germination (***Sreenivasulu et al., 2008***; ***Shutov and Vaintraub, 1987***; ***Hammerton and Ho, 1986***; ***Dal Degan et al., 1994***). Similarly, just prior to radicle sprouting, genes encoding proteins involved in amino acid biosynthesis and transport are upregulated, whereas genes involved in amino acid degradation are largely unresponsive (***Ma et al., 2017***). This suggests that nitrate reductase activity is not required to reassimilate nitrogen from amino acid catabolism. Our results suggest that after feeding enriched amino acids, the spread of the tracer across all proteinogenic soluble amino acid pools is rather limited and only increases at low sub-optimal temperature. Thus, it appears that amino acid degradation and reassimilation are suppressed processes during barley germination at optimized temperature. Instead, barley seeds may be genetically tuned to rely on amino acid mobilization during the early stages of embryo development. Therefore, using labelled amino acids to introduce the tracer into germinating barley seedlings may be the best and only strategy to follow *in vivo* isotopic fluxes into protein.

Germinating barley seedlings have at least four systems for successful amino acid uptake (***Salmenkallio and Sopanen, 1989***), all of which depend on storage proteins or their hydrolysis products being taken up into the scutellum for further hydrolysis and utilization (***Higgins and Payne, 1981***). Thus, the supply of ^15^N-labeled amino acid compounds in the germination media ensures that these nitrogen atoms are introduced into and utilized by the plants. However, because of the generous availability of endogenous amino acid resources, the incorporated ^15^N-labeled amino acids are diluted in the roots of germinating seedlings. Moreover, at low temperatures, all enzymatic activities and cellular dynamics slow down. Therefore, mobilization of amino acids and peptides for nitrogen supply during germination is also likely to be affected by lower temperatures. Our results suggest that barley seedlings subjected to acclimation take up more labelled amino acids through their roots and use / spread their nitrogen supply across soluble amino acid pools. Thus, it is possible that nitrogen deficiency resulting from slowed nutrient mobilization is compensated for by increased uptake of nutrients through the roots, and since amino acid and peptide transporters are already available from the germination process, these compounds would be adsorbed. The bottom line is that the physiological transition triggered by acclimation to low temperatures causes differential incorporation of amino acids carrying the tracer, differential spread of the tracer across amino acid pools and different accumulation of soluble amino acids, and all of these differences must be accounted for if biological insights are to be derived from tracking the tracer incorporation into polypeptides.

### Ribozyme-mediated ^15^N Incorporation Into Protein

In all living organisms, the proteome is synthesised by ribosomes, whose ribozyme activities catalyse the formation of peptide bonds between the existing peptidyl-tRNA and the subsequent aminoacyl-tRNA (***Rodnina, 2013***). Thus, the natural pathway of ^15^N-labeled amino acids is to conjugate with tRNAs and be transported to ribosomes, where they enter the elongation cycle and end up as a monomer in a synthesised polypeptide. Amino acids exist as soluble pools in the cytoplasm and are loaded onto aminoacyl-tRNAs, which are present in much lower proportions as compared to amino acid pools and are much more labile, i.e., their turnover is extremely rapid (***Gomez and Ibba, 2020***), suggesting that the enrichment in soluble amino acid pools is a valid proxy for isotope enrichment in aminoacyl-tRNA conjugates.

The main activity of ribosomes is autocatalysis (***Reuveni et al., 2017***), and as such the tracer is expected to be incorporated first into the machinery within one degree of ribosomes (***Bowman et al., 2020***), which include rProteins, aminoacyl-tRNA synthetases, tRNAs, initiation, elongation and release factors. The cellular proteome within one degree of ribosomes is part of macromolecular complexes. Thus, by purifying a complexome proteome, one can recover translation-related multiprotein complexes in a near-native state while co-purifying other complexes. In this way, it is possible to test the link between ribosome structural divergence and altered rates of protein synthesis at sub-optimal low temperature. Altered protein synthesis can be caused by direct or indirect translational control determining which transcripts are translated under the limited growth of cold acclimation. Direct translational control implies an altered and selective ribozyme-functionality shaping the proteome. In our system, translation during cold is carried out by a ribosomal population that is heterogeneous and substoichiometric in its rProtein composition. Altered rProteome compositions confer metazoan ribosomes the ability to selectively recruit transcripts for translation, i.e., to specialize (***Shi et al., 2017***; ***Genuth and Barna, 2018***). Additionally, Plants have an increased number of rProtein paralogs compared to all higher metazoans (***Barakat et al., 2001***), and the fate of duplicated genes usually leads to novel and divergent functions (***Kosová et al., 2021***). Thus, it is probable that specialized rProtein proteoforms may equip heterogenous ribosomes to perform direct translational control, efficiently adapting them to cold temperatures.

### Translational Dynamics of Heterogeneous Ribosomes

In Arabidopsis, cold heterogeneous translating ribosomes exhibit rProtein substoichiometry around the polypeptide exit tunnel (PET), with many of the rProteins being relatively removed during cold (***Martinez-Seidel et al., 2021a***). Here, we report that barley ribosomes also exhibit, on average, subtractive heterogeneity (***Briggs and Dinman, 2017***) in the cold-ribosomal population around protein uL4 and uL29, the former being essential for PET assembly during ribosome biogenesis (***Lawrence et al., 2016***; ***Gamalinda and Woolford, 2014***; ***Stelter et al., 2015***; ***Pillet et al., 2015***), as its internal loops form the constriction sites of the nascent PET (***Micic et al., 2022***) along uL29. The PET and its assembly are likely to be particularly critical during cold acclimation because both yeast (***Hung and Johnson, 2006***) and plants (***Schmidt et al., 2013***) have a 60S maturation factor that when knocked out, leads to cold sensitivity, namely Rei-1 in yeast and its homolog REIL in plants. The functional role of Rei-1 in yeast is to insert its C-terminus into the PET to check the integrity of the tunnel as a quality control step before making 60S subunits translationally competent (***Greber et al., 2016***). Thus, subtractive rProtein heterogeneity near the tunnel could indicate higher rRNA disorder, defective tunnel assembly, and/or defective structure, causing the observed need for PET quality control during cold.

On the other hand, the 60S rProteins accumulated in the cold population of ribosomes, uL18 and eL30, are located near important intersubunit bridges (***Martinez-Seidel et al., 2021b***; ***Tamm et al., 2019***). Similarly, the population of 40S subunits, which shows accumulation of specific rProteins only during cold, has two rProteins, eS6 and eS1, that also form important intersubunit bridges in the form of connections between the 40S and 60S and their constituent rProteins (***Martinez-Seidel et al., 2021b***; ***Tamm et al., 2019***). The third rProtein more abundant in 40S subunits, eS10, links the large uS3 hub (containing the ribosomal region adjacent to the tRNA-mRNA entry sites) to the uS13-uL11 subunit bridge (***Martinez-Seidel et al., 2021b***). Bridges between subunits in bacteria have been shown to directly affect initiation factor-dependent translation (***Kipper et al., 2009***). Thus, these observations suggest that the rProteins accumulated during cold in ribosomal populations are related to subunit connectivity and may influence their association during initiation, elongation or termination.

### Translation Initiation: Newly Synthesised Complexes

Complexes related to translation initiation were accumulated during cold acclimation in our system due to synthesis of their protein components, implying tight control over what type of translation initiation complexes form and participate in 40S transcript association. Translation initiation is a sequential and complex process that is highly conserved (***Jackson et al., 2010***; ***Hashem and Frank, 2018***) and nonetheless exhibits peculiarities in its regulation in plants (***Castellano and Merchante, 2021***). Translation initiation begins with the binding of multiple factors to the 40S subunit to form a 43S pre-initiation complex (PIC) (***Aylett et al., 2015***; ***Hashem et al., 2013***; ***Majumdar et al., 2003***), which then binds the mRNA to be translated, supported by multiple factors, to form the 48S initiation complex (IC) (***Brito Querido et al., 2020***; ***Pisareva and Pisarev, 2014***). Finally, the IC is supported by several factors to allow the 60S subunits to connect, making the elongation process competent in the newly formed 80S monosome (***Fringer et al., 2007***; ***Shin et al., 2002***). In our study, we found that several protein components of the initiation machinery are both significantly accumulated and synthesised during cold:

1. First, the eukaryotic translation initiation factor 3 subunits A, B, C, and E (eIF3A-C,E). The eIF3 complex consists of 13 subunits (A-M) and is the most complex initiation factor in eukaryotes and also the largest (***Des Georges et al., 2015***; ***Zhou et al., 2008***). Moreover, eIF3 has been associated with various pathological conditions in higher metazoans (***Gomes-Duarte et al., 2018***). Subunits A and C bind eIF3 to 40S subunits via the platform on the solvent side (***Aylett et al., 2015***) and can also interact with eIF1 and eIF4G via subunit E (***LeFebvre et al., 2006***). Thus, 3/4 of the preferentially accumulated and synthesised eIF3 subunits serve as anchors between ribosomes and mRNA recruitment factors. Subunit B is part of the eIF3b-i-g module and presumably interacts with the 40S subunit directly at the mRNA entry channel by occupying it (***Chiu et al., 2010***). Accumulation of these specific subunits of the eIF3 complex is associated with cancer in humans via increased (***Scoles et al., 2006***; ***Xu et al., 2012***; ***Wang et al., 2013***) and selective (***Dong et al., 2004***; ***Parasuraman et al., 2017***) translational output.
2. Second, eukaryotic translation initiation factor 2 subunit 1 (eIF2*α*). This eIF catalyses the first step of 40S - initiator-tRNA (Met-tRNA) association (***Hinnebusch, 2017***). This factor is the central element of the integrated stress response in eukaryotes (***Pakos-Zebrucka et al., 2016***), as it leads to a global decrease in protein synthesis through a phosphorylation event mediated by eIF2*α* kinases, while promoting the selective translation of specific transcripts whose protein products are required for survival (***Lu et al., 2004***). In this case plants may be using this robust and well-characterized stress response for successful acclimation to sub-optimal low temperatures.

Importantly, the significant accumulation of a protein fraction enriched in rProteins during cold, which we reported previously (***Martinez-Seidel et al., 2021c***) and demonstrated again in this study (Figure 5 - Figure S1), originated from complexes related to ribosome biogenesis and translation initiation, highlighting the functional importance of translation initiation during cold acclimation and assigning a potential regulatory and functional role to the group of cold heterogeneous rProteins. The accumulation of machinery at both sides of competent ribosomes, i.e. pre-ribosomes and translation initiation complexes, along with the mantained ratio of 40S to 60S subunits suggests that the limiting step during cold acclimation is successful initiation. Thus, the set of competent 60S subunits assembled during cold might select which transcript to translate based on an excess of PICs and ICs.

### Ribosome Biogenesis: Assembled and Remodelled Ribosomes

The increasing complexity uncovered in the process of ribosome biogenesis suggests that the highly energy demanding process of assembling ribosomes is connected to major environmental and developmental cellular responses (***Lindahl, 2022***). In our study, ribosome biogenesis complexes accumulated during cold acclimation due to the lack of degradation, as their protein components did not take up the ^15^N tracer and yet were significantly more abundant during cold compared with control conditions. The only pre-ribosomal complex that accumulated newly synthesised protein components was the small subunit processome. The small subunit processome is the earliest pre-40S complex found to date in eukaryotes, and it uses many accessory factors to process and mature 40S subunits (***Barandun et al., 2017***). Therefore, the protein components that are newly synthesised during cold might indicate alternative processing of pre-40S subunits, which could be related to the reported cold-specific accumulation of 40S rProteins in assembled SSU complexes. At the structural rProtein level, there were fewer competent ribosomes during cold, and only 60S subunit components were generally and preferentially synthesised during cold, to such an extent that GO enrichment of the 60S structural component category was detected when the cold-synthesised proteome was examined. Thus, our results suggest that there is restructuring due to altered LSU rProtein synthesis in cold-acclimated 60S large subunits and variability in processing of pre-40S small subunits of cytosolic ribosomes due to differential protein synthesis of components from the small subunit processome. Both aspects are complemented by increased abundances of ribosome biogenesis complexes, suggesting that ribosome heterogeneity arises in part from the ribosome assembly line at low sub-optimal temperatures.

At the individual protein level, it is clear that ribosomes are assembled or remodelled using new and old rProteins. Only a subset of rProteins significantly incorporated the tracer ^15^N, confirming the previously observed fact that in plants cytosolic rProteins exhibit the greatest variability in degradation rates among protein components of large cellular complexes (***Li et al., 2017***). This variability makes economic sense, as ribosome assembly and protein biosynthesis impose the greatest cellular costs (***Shore and Albert, 2022***) and continued synthesis of each component within the translational apparatus would be detrimental. In addition, rProteins in plants have a half-life of about 4 days (***Salih et al., 2020***), and given the slower cell dynamics at cold temperatures, relying solely on ribosome biogenesis to control translational dynamics would be too slow. Therefore, remodelling old ribosomes to adapt them for the work at hand could be an efficient way to alter their function. For example, in higher metazoans, ribosomes in neuropil (far from the nucleolus) are remodelled *in situ* rather than undergoing a new cycle of ribosome biogenesis and subsequent transport to dendrites and axons, and this is a means of regulating local protein synthesis depending on the cellular context (***Fusco et al., 2021***). Whether this is the case in plants remains to be tested, because we do not report rRNA synthesis rates required to determine whether a complete biogenesis cycle generated the heterogeneous ribosomes using new and old rProteins or whether, on the contrary, these ribosomes were remodelled in the cytoplasm.

### Translational Outcome of Heterogenous Ribosomes: A Proteome Shift

The assembled and remodelled heterogeneous ribosomes are able to cause a significant part of the proteome shift observed during cold (the rest of the shift is due to the lack of degradation of specific proteins). In addition to complexes related to translation, cold-remodelled ribosomes significantly affect the synthesis and accumulation of protein folding machinery, complexes from the endoplasmic reticulum (ER) and Golgi, nuclear complexes and complexes related to the cell cycle, cell wall and microtubule complexes, and protein complexes from the outer mitochondrial membrane. All of these complexes and their individual proteins could be of great importance for acclimation, as the plant accumulates them beyond the level reached in plants grown at optimal temperature, despite its limited resources and slow growth dynamics.

We found that 6/8 subunits of the cytosolic chaperonin T-complex-protein 1 ring complex or chaperonin-containing TCP-1 (CCT) are preferentially synthesised during cold, leading to accumulation of the complex, which may be directly related to the altered functionality of cold heterogeneous ribosomes. The CCT binds and promotes protein folding of newly synthesised polypeptides (***Lopez et al., 2015***; ***Yébenes et al., 2011***) or promotes their aggregation and thus protein degradation to maintain proteostasis (***Lopez et al., 2015***; ***Hartl et al., 2011***; ***Spiess et al., 2004***). We have shown that during cold in plants, ribosome remodeling, i.e., subtractive heterogeneity, occurs in the proteome surrounding the PET, and the signature of a defective tunnel is protein misfolding (***Micic et al., 2022***; ***Peterson et al., 2010***; ***Wruck et al., 2021***). Thus, it is conceivable that an altered tunnel structure leads to increased protein misfolding and this forces the plant cell to produce more CCT complexes to properly fold or aggregate misfolded proteins in order to fix or degrade them, respectively. Moreover, we found that several heat shock proteins are preferentially synthesised and accumulated at cold temperatures, underscoring that protein misfolding is an urgent problem that plants need to address during acclimation to sub-optimal low temperatures.

We also found that nuclear and cell cycle complexes were preferentially synthesised and accumulated. Among them was condensin, which in addition to its role in the cell cycle promoting chromosome assembly, is also known to directly regulate gene expression (***Iwasaki et al., 2019***; ***Li et al., 2015***). Plants stop their mitotic activity in response to cold (***Ashraf and Rahman, 2019***), and thus condensin is less likely to accumulate to promote further cell cycle progression, but rather may accumulate to promote mechanisms of transcriptional control in response to cold.

All other preferentially synthesised and accumulated complexes can be classified into the machinery that transports and targets the cell membrane and wall. Among this type of complexes we found ER and Golgi components as well as cell wall and microtubule proteins. They were all directly related to protein transporters, glycosyltransferases, membrane transport components, and cell wall structure. The integrity of cell membranes and walls is threatened in plants during cold as membrane fluidity decreases and dehydration increases (***Takahashi et al., 2020***). Therefore, physical changes such as remodelling of the lipidome of membranes (***Barrero-Sicilia et al., 2017***) are essential to survive cold (***Johnson, 2018***). For example, abundance changes of specific lipid species in other cereal models confer cold resistance (***Cheong et al., 2022***). Such membrane remodelling mechanisms in plants can rely on enzymatic activity to deliver new components needed to enhance cold acclimation (***Fourrier et al., 2008***). Many of the proteins that are transported to the cell periphery are initially synthesised near the ER - Golgi organelles, where the shuttles that transport them are ready to deliver them to their site of function. We have provided evidence that the transport machinery is active during cold, accumulating and being newly synthesised. This may be part of the plant’s attempt to mitigate the loss of membrane integrity by transporting necessary membrane and wall components or the enzymes to produce them. Most importantly, translation of this sub-proteome demonstrates that cold-heterogeneous ribosomes are able to directly or indirectly control their translational output to efficiently acclimate.

Cold-heterogenous ribosomes also synthesise components of other macromolecular complexes without accumulating them. This type of synthesis implies potential remodelling of complexes that continue to be present in equal amounts due to ongoing protein turnover. Complexes that fall into this category include the 60S ribosmal subunit, translation initiation, ER-Golgi, proteasome, vacuolar proton-transporting ATPase, and the oxoglutarate metabolon.

## Conclusions

With careful consideration of the ^15^N isotope flux and plant phenotype, we were able to monitor tracer incorporation into digested peptides of proteins at the complexome proteome level and compare between experimental conditions. Our strategy can be applied to any system that transitions between different biological steady states to study the dynamics of protein synthesis, as long as the right variables can be measured. We have made available our equations and complete bioinformatics method as a public R package, i.e., the ProtSynthesis R package. We applied this strategy to understand the transition of proliferative root tissue from germinating barley seedlings to a cold acclimated state. The proliferating root tissue of germinating barley seedlings undergoing cold acclimation, like Arabidopsis, requires ribosome biogenesis to overcome the initial stimulus. In addition, plants build remodelled and heterogeneous ribosomes that cause a shift in the proteome. To characterize the heterogeneity, we mapped the relative stoichiometry of ribosome-assembled rProteins and their synthesis rates using proteome-wide ^15^N labelling to determine which part of the rProteome shift is due to synthesis and which part is due to reuse of pre-existing rProteins. We can currently conclude that plants significantly modulate the relative synthesis rates of ribosomebound rProteins differentially when confronted with environmental factors, such as a shift to suboptimal temperature, and that such modulation appears to be independent of *de novo* ribosome assembly. Moreover, ribosomes remodelled in the cold exhibit subtractive heterogeneity around the polypeptide exit tunnel (PET) and an accumulation of specific rProteins, in both 40S and 60S subunits, that are structurally linked to key intersubunit bridges. In addition, we examined general proteome shifts and found that 43S and 48S translation initiation complexes are preferentially synthesised and accumulate during cold, leading to a higher requirement for 60S subunits, which are at a constant ratio with 40S subunits but appear to be insufficient to form elongation-competent 80S monosomes and solve the over-accumulation of initiation complexes. We therefore hypothesize that 60S subunits are not able to bind all of the translation initiation complexes, and consequently they selectively associate with specific transcript-associated 48S complexes. This hypothesis is supported by the cold-induced heterogeneity, which mainly relates to the association of 40S and 60S subunits and as such could be a way to identify translational needs inherent to the cold context. The other major shift in the newly synthesised proteome is a response to protein aggregation and misfolding, which we propose is linked to missing rProteins around the PET in cold-remodelled ribosomes. This mechanism may be a second layer of translational control that allows ribosomes to misfold and target for degradation the part of the proteome that is currently not needed. From this study, we can currently conclude that there are major responses in the plant translational apparatus during cold that cause ribosomes to build a proteome to respond to the consequences of their own structural adaptations. Concomitantly the cold-heterogeneous ribosomes are able to directly or indirectly cause proteome shifts to remodel the cellular membrane and cell wall as part of the agenda to transition to an acclimated state and eventually resume growth.

## Methods and Materials

### Plant rearing

#### Surface Seed Sterilisation and Imbibition

*Hordeum vulgare* cultivar Keel seeds were obtained from The University of Melbourne from previous studies (***Gupta et al., 2019***). Seeds were placed inside sterile 50 mL falcon tubes (max 2 g per falcon tube approx. 40 seeds) amounting to a total of approx. 600 seeds (i.e., 15 falcon tubes). The non-biological materials were surface sterilised with 70 % ethanol and placed inside a clean bench, followed by UV-sterilisation. The seeds were soaked in 70 % ethanol and shaken gently for 1 min, ethanol was then discarded. A 1 % bleach solution (0.042 % sodium hypochlorite) was then added to the falcon tube and gently shaken for 10 min, after which the bleach solution was discarded. The seeds were rinsed five times with sterile MilliQ H_2_O and gently shaken for 5 min each time to completely remove the hypochlorite. The water was discarded. After straining the water, the seeds were soaked in sterile MilliQ H_2_O and the falcon tubes were wrapped in aluminium foil to prevent any light exposure for 14 to 18 hours, half of which was at 25 °C and the other half at 18 °C, to mimic the daily temperature fluctuations and initiate imbibition of the seeds.

#### Seedling Germination and Treatment

Seeds were germinated and treated in complete perceived darkness by using a green light filter to cover the light entering the clean bench when the seeds were transferred to plates. For germination, seeds were transferred to 72 Petri dishes that were filled with 10 ml non-labelled, non-supplemented Scheible medium (***Scheible et al., 2004***) for 48 hours, 8 to 16 seeds were transferred into each petri dish. The dishes were sealed with micropore tape, wrapped in aluminium foil and placed in a phytotron growth chamber (Weiss Technik, Germany) with temperature settings of 25 °C for 16 h and 18 °C for 8 h until the completion of 48 h to allow germination. When 48 hours had passed, six germinated plants from a petri dish were harvested and processed to calculate RGR at time point zero (to be explained below). Then, working on the clean bench and under a green light filter, the medium in the dishes was disposed and exchanged for labelled and non-labelled supplemented media. 42 dishes with 10 mL non-labelled Scheible medium supplemented with 0.5 mM ^14^N Serine and Glycine and 30 dishes with 10 mL labelled Scheible medium supplemented with 0.5 mM ^15^N Serine (609005, Lot: MBBB0411V, Sigma Aldrich) and Glycine (299294, Lot: MBBC7772, CAS: 7299-33-4, Sigma Aldrich). Half of the labelled and non-labelled dishes were shifted to 4 °C to induce cold acclimation for 5 days. The other half remained in the control growth chamber with temperature fluctuation of 25 °C for 16 h and 18 °C for 8 h.

### Plant Harvest and Phenotyping

For phenotyping, each individual plant was considered a biological replicate. In total 12 dishes were used, one per treatment, per time-point. Six cold-germinated and six control-germinated plants were harvested each day from day 0 until day 5. Immediately, the seedlings were scanned in order to determine the length, width, volume and area of the roots. Then, the roots of each plant were harvested separately and the excess media was completely removed with a paper towel. Samples were wrapped in a folded piece of Pergamin paper and immediately weighed for the fresh weight (FW), and then dried for 70 h at approx. 70 °C and weighed again for dry weight (DW). For all subsequent analysis, root tips were collected in 1.5 cm segments on the fifth day of acclimation. Labelled and non-labelled plants reared at cold and control temperatures were collected for primary metabolome and complexome / ribo-proteomic assays. Root segments of plants from the remaining 60 dishes were harvested by pooling root segments from 5 dishes per biological replicate (for a total of 3 biological replicates per labelling - temperature treatment combination). Briefly, 1.5 cm segments of the root tip were collected using a sharp blade and flash frozen with liquid nitrogen. Handling the plant material in frozen conditions, root pools were grinded in a prefrozen mortar and pestle, followed by preparation of 200 mg and 60 mg aliquots for complexome / ribo-proteomic and primary metabolome analyses respectively. Ground plant material was stored at -80°C until further use.

### Morphometric Image Processing

Images of the complete roots were acquired with an image analysis system and scanner (Perfection V800 Photo, Epson). Subsequently the software winRHIZO (Regent Instruments Inc., version released as 2019a) was used to delineate the root tissue and quantify the relevant morphological variables, root length, root average diameter, root volume, number of forks or bifurcations, number of tips, root length per volume. The software parameters were: ImgType - Grey, CalibMeth - Intr, TPU Units - cm, PxSizeH - 0.006353, PxSizeV - 0.006347 [CalFile] - Scanner.Cal, PxClassif - GreyThdAutom-57, Filters - Smooth Off Area Off LWRatio Off, Fractal PxMin PxMax - Off.

### Primary Metabolome Analysis

To extract metabolites, 360 μL of pre-cooled extraction mix containing Methanol:Chloroform:Water (2.5:1:1(v/v)) and 30 μL of U-^13^C sorbitol (0.2 mg/mL), as internal standard, was added to 60 mg flash-frozen, grounded root tip tissue, vortexed vigorously and incubated at 70 °C for 15 min. Once the samples were cooled down to room temperature, 200 μL of CHCl_3_ was added and incubated at 37 °C for 5 min with shaking. Phase separation was induced by adding 400 μL H_2_O, vortexed and centrifuged at 20,800 rcf for 10 min. 160 μL aliquot from the upper polar phase was transferred to fresh 2 mL microvials and dried by vacuum centrifugation for 18 hours at room temperature. The dried samples were stored at -20°C until further use.

Primary metabolites were analysed by gas chromatography-mass spectrometry (GC-MS) of methoxyaminated and trimethylsilylated metabolite preparations (***Erban et al., 2020***). Metabolite extraction, chemical derivatization were as previously described. C_10_, C_12_, C_15_, C_18_, C_19_, C_22_, C_28_, C_32_, C_36_ n-alkane mixture was added to each sample for retention index calculation. Samples were processed using a Factor Four Capillary Column VF-5ms of dimensions 30 m length, 0.25 mm internal diameter and 0.25 mm film thickness (Variant Agilent) mounted to an Agilent 6890N gas chromatograph with split/splitless injector and electronic pressure control up to 150 psi (Agilent, Böblingen, Germany). Mass spectrometric data were acquired through a Pegasus III time-of-flight mass spectrometer (LECO Instrumente GmbH, Mönchengladbach, Germany), and in parallel using the same samples at high mass resolution using a micrOTOF-Q II hybrid quadrupole time-of-flight mass spectrometer (Bruker Daltonics, Bremen, Germany) with a multipurpose APCI source. Detailed GC-electron spray ionization TOF-MS settings were as reported previously (***Erban et al., 2020***).

Metabolites were annotated and identified by mass spectral and retention index matching to data of authenticated reference compounds from the Golm Metabolome Database (***Kopka et al., 2005***).

### Ribosome Enriched Proteomics

#### Protease Considerations

Lys-C over Trypsin: Amino acids enriched in RNA protein-binding domains are histidine, arginine and lysine, which are all basic amino acids and. The rProteome is enriched in basic amino acids in order to be able to bind rRNA. Trypsin cleaves peptide sequences at the C-terminal of lysine and arginine residues, thus it would digest rProteins into smaller pieces as compare to Lys-C, which cleaves peptide sequences only at the C-terminal side of lysine residues. Thus Lys-C cuts sets of basic proteins such as the rProteins into significantly longer pieces as compared to trypsin (Figure 6 - Figure S1).

#### Ribosomal Protein Purification and Processing

Cell lysis was induced in grounded plant tissue using reported methods (***Firmino et al., 2020***; ***Martinez-Seidel et al., 2021c***) with minor modifications. Briefly, aliquots were placed in liquid nitrogen-cooled mortars and mass spectrometry friendly ribosome extraction buffer (MS_*f*_ -REB) was added at a buffer (V) to tissue (FW) ratio of two. The extract was then homogenised for 20 minutes while the mortars stayed on ice to prevent temperature from rising. Big particles were filtered through a pre-made, autoclaved and tip-amputated 5 mL pipette tip containing a ©Miracloth clog inside, and the filtrate was aliquoted in 2 mL microcentrifuge tubes. Samples were centrifuged at 14,000 x g for 20 min (4°C) to pellet insoluble cell debris and supernatants were transferred to violet QIAshredder mini spin columns (Qiagen, Australia) and centrifuged again for one minute. Sample volume was adjusted to 4.5 ml in order to fill the ultracentrifuge tubes until 50 %. Subsequently, extracts were loaded carefully into thick-walled polycarbonate tubes with three-piece caps (10.4 mL, Polycarbonate Bottle with Cap Assembly, 16 × 76 mm - 6Pk, 355603, Beckman Coulter, USA) pre-filled with 2.5 mL sucrose cushion (SC) solution.

MS_*f*_ -REB: 0.2 M of Tris, pH 9.0, 0.2 M of KCl, 0.025 M of EGTA, pH 8.0, 0.035 M of MgCl_2_, 1 % (W/V) octyl beta-D-glucopyranoside (98 %, O8001, Sigma Aldrich, Australia), 0.18 mM cyclohexamide (Sigma Aldrich, Australia), 5 mM Dithiothreitol (R0861, Thermo Fisher, Australia), 1 mM Phenylmethylsulfonyl fluoride (36978, Thermo Fisher, Australia), 1X protease inhibitor cocktail (cat. No. P9599, Sigma Adrich, Australia).

SC: 0.4 M of Tris, pH 9.0, 0.2 M of KCl, 0.005 M of EGTA, pH 8.0, 0.035 M of MgCl_2_ × 6H_2_O, and 60 % sucrose (Molecular Biology Grade, 573113, Sigma Aldrich, Australia), 0.18 mM cyclohexamide (Sigma Aldrich, Australia), 5 mM Dithiothreitol (R0861, Thermo Fisher, Australia), 1 mM Phenylmethylsulfonyl fluoride (36978, Thermo Fisher, Australia), 1X protease inhibitor cocktail (cat. No. P9599, Sigma Adrich, Australia).

Loaded samples were centrifuged at 4°C, 330,000 x g / 60,000 RPM for 4.5 hours using a TY 70.1Ti rotor (Type 70.1 Ti Rotor, Beckman Coulter, USA) loaded into an Optima XE-100 Ultracentrifuge (Beckman Coulter, USA). After centrifugation, the supernatant was removed including the sucrose cushion taking care that the only solution in contact with the pellet was the SC. Tubes were completely dried by placing them upside down for a couple of minutes and dried pellets stored at -80°C until further usage. Ribosomal pellets were resuspended in 60 μL, freshly prepared, GuHCl to dissociate rProteins and TFA was added to 1 % final volume to induce precipitation of nucleic acids. The solution was then centrifuged in a microcentrifuge at 20,800 x g for 20 minutes and the supernatant recovered. Protein content was determined in samples using the bicinchoninic acid (BCA) kit (Thermo Scientific, United States) assay. As a control *Escherichia coli* ribosomes (P0763S, NEB, Australia) were used in 4 μL aliquots (approx. 2000 A260 units that are equivalent to 102 μg of ribosomes and 23 μg of rProtein) to undergo the full protocol (Figure 6 - Figure S2), confirming the integrity of ribosomal complexes when passing through the SCs solution and subsequent rProtein dissociation.

Protein amounts were standardised to the minimum concentration, i.e., 13.7 μg in 50 μL 6M GuHCl, 1 % TFA. Proteins were reduced and alkylated by adding tris(2-carboxyethyl)phosphine (TCEP) (77720, Thermo Scientific, United States) and iodoacetamide (IAA) (A3221, Sigma Aldrich, Australia) to 10 mM and 55 mM respectively, and shaking for 45 minutes at 37°C. The alkylation step was shaken in the dark. Acetonenitrile was then added to 70 % and a 10:1 ratio of magnetic beads (Hydrophilic-Part no: 45152105050250, GE Healthcare plus Hydrophobic-Part no: 65152105050250, GE Healthcare, Australia) were added and mixed with the solution. Beads were prepared according to the manufacturers instructions to a concentration of 20 *μ*g/*μ*L stock. Solution sat for 20 minutes with two pipette mixes, one every 10 minutes. Tubes were placed on a magnetic rack (DynaMag-2, 12321D, Life Technologies) and allowed to separate for 30 seconds. Washes were performed while the tubes remained in the rack. 1 ml of neat acetonitrile was added for 10 seconds and removed followed by 1 ml of 70 % ethanol for 10 seconds and removed. Tubes were removed from the rack and 1:10 protein (μg) to digestion buffer (μL) was immediately added. The digestion buffer (25mM triethylammonium bicarbonate (TEAB)) contained the Lys-C protease (P8109S, NEB, Australia) at a 1:20 protease to protein ratio. Samples were incubated for 18 hours at 37°C at 1000 RPM in a thermomixer (Eppendorf, Australia). TFA was added to 1 % to quench the reaction and then the tubes were put on the magnetic rack and the supernatant transferred to new tubes, twice. Finally a centrifugation step at 14,000 x g was performed to get rid of any residual beads and only 90 % of the supernatant was recovered. The recovered fraction was frozen for an hour at -80°C and then freeze-dried. Peptides were resuspended in MS-loading buffer (2 % Acn + 0.05 % TFA) and loaded into a LC-MS/MS platform.

#### LC-MS/MS Analysis

All samples were analysed in the Nano-ESI-LC-MS/MS. The Nano-LC system, Ultimate 3000 RSLC (Thermofisher Scientific, San Jose, CA, USA) was set up with an Acclaim Pepmap RSLC analytical column (C18, 100 Å, 75 μm × 50 cm, Thermo Fisher Scientific, San Jose, CA, USA) and Acclaim Pepmap nano-trap column (75 μm × 2 cm, C18, 100 Å) and controlled at 50 °C. Solvent A was 0.1 % v/v formic acid and 5 % v/v dimethyl sulfoxide (DMSO) in water and solvent B was 0.1 % v/v formic acid and 5 % DMSO in ACN. The trap column was loaded with digested peptides at an isocratic flow 3 % ACN containing 0.05 % trifluoroacetic acid (TFA) at 6 μl/min for 6 min, followed by the switch of the trap column as parallel to the analytical column.

To measure peptides from barley experimental rProteome samples, the gradient settings for the LC runs, at a flow rate 300 nl/min, were as follows: solvent B 3 % to 23 % in 89 min, 23 % to 40 % in 10 min, 40 % to 80 % in 5 min, maintained at 80 % for 5 min before dropping to 3 % in 0.1 min and equilibration at 3 % solvent B for 9.9 min. An Exploris 480 Orbitrap mass spectrometer (Thermo Fisher Scientific, San Jose, CA, USA) with nano electrospray ionization (ESI) source at positive mode was employed to execute the MS experiments using settings of spray voltages, ion funnel RF, and capillary temperature level at 1.9 kV, 40 %, 275 °C, respectively. The mass spectrometry data was acquired with a 3-s cycle time for one full scan MS spectrum and as many data dependent higher-energy C-trap dissociation (HCD)-MS/MS spectra as possible. Full scan MS spectra feature ions at m/z of 300-1600, a maximum ion trapping time of 25 msec, an auto gain control target value of 3e^6^, and a resolution of 120,000 at m/z 200. An m/z isolation window of 1.2, an auto gain control target value of 7.5e^4^, a 30 % normalized collision energy, a first mass at m/z of 120, an automatic maximum ion trapping time, and a resolution of 15,000 at m/z 200 were used to perform data dependent HCD-MS/MS of precursor ions (charge states from 2 to 6).

To measure peptides from commercially available *Escherichia coli* ribosomes processed into rProteome samples gradient settings for the LC runs, at a flow rate 300 nl/min, were as follows: solvent B 3 % to 23 % in 59 min, 23 % to 40 % in 10 min, 40 % to 80 % in 5 min, maintained at 80 % for 5 min before dropping to 3 % in 0.1 min and equilibration at 3 % solvent B for 9.9 min. An Eclipse Orbitrap mass spectrometer (Thermo Fisher Scientific, San Jose, CA, USA) with nano electrospray ionization (ESI) source at positive mode was employed to execute the MS experiments using settings of spray voltages, ion funnel RF, and capillary temperature level at 1.9 kV, 30 %, 275 °C, respectively. The mass spectrometry data was acquired with a 3-s cycle time for one full scan MS spectra and as many data dependent higher-energy C-trap dissociation (HCD)-MS/MS spectra as possible. Full scan MS spectra features ions at m/z of 375-1500, a maximum ion trapping time of 50 msec, an auto gain control target value of 4e^5^, and a resolution of 120,000 at m/z 200. An m/z isolation window of 1.6, an auto gain control target value of 5e^4^, a 30 % normalized collision energy, a maximum ion trapping time of 22 msec, and a resolution of 15,000 at m/z 200 were used to perform data dependent HCD-MS/MS of precursor ions (charge states from 2 to 6).

Complete dataset proteomics submissions have been deposited to the ProteomeXchange Consortium (***Deutsch et al., 2020***) via the PRIDE (***Perez-Riverol et al., 2019***) partner repository with the dataset identifiers PXD032923 for *H. vulgare* experimental samples (**DOI**: 10.6019/PXD032923) and PXD032938 for *E*.*coli* control samples (**DOI**: 10.6019/PXD032938).

### Data Analyses

#### Phenotyping

Relative growth rates using fresh, dry weight and varied phenotype measurements were calculated using R programming language with equation 1. The output units were mg * (mg^−1^ * h^−1^). W is total weight accumulation at t_*f*_ and it is used in equation 1 to transform the growth rates into fractional of the final weight. Delta W or *dW* is the change in fresh or dry weight accumulation in milligrams from time 0 or non-germinated until delta t or *dt*, which represents hours after germination. Thus assuming an initial root weight of 0 leads to *dW*_*t*_ being represented by the weight measurement at time-point t. The Tukey-HSD test was performed after an ANOVA with a confidence level of 95 %.

#### Primary Metabolome

Amino acid abundances were analytically derived from GC-EI-ToF-MS acquired data using the software TagFinder (***Luedemann et al., 2008***) and the Golm Metabolome Database (***Kopka et al., 2005***). Extraction, standardisation, derivatization and GC-MS analytics were performed according to Erban et al., 2020 (***Erban et al., 2020***). Three biological replicates were measured in technical triplicates. For high abundant metabolites reaching the detection limit measurements of all samples were repeated with a 1:30 dilution (split) of extracts. Compounds were manually annotated in TagFinder and representative Tags for each metabolite chosen. Metabolome data were normalized to the levels of an internal ^13^*C*_6_ sorbitol standard (CAS 121067-66-1), the background levels of the blanks were subtracted and data were normalized to the fresh weight of plant material in each sample. For Table S2, primary metabolome data were analysed with the “OmicsUnivariateStats.R” function of the RandoDiStats R package. Here, missing values were replaced by a small normally distributed numeric vector. Additionally, the fold change of metabolite abundances under cold conditions as well as the logical induction of metabolites (absence-presence-scenarios) were calculated.

^15^N enrichment percentages of labelled metabolite pools was analytically derived from a multiplexed GC-EI-ToF-MS and GC-APCI-qToF-MS platform. In the first case, the workflow entailed baseline correction of the raw chromatogram files using the vendor software and transformation into CDF files. Pre-processing of the chromatograms for increasing data matrices quality (internal standard normalization and chromatogram alignment, mass scan width synchronisation) using TagFinder. Similarly, in the latter case, the vendor software was used to find the amino acid peaks in the chromatograms. peaks were manually mined and integrated in order to derive relative abundances. Every step of the targeted manual annotation of N containing mass tags is deposited in Table S3. Since ^15^N feeding can cause differential abundances in monoisotopic fragments from the same amino acid analyte depending on the lack or presence of a N atom, multiple fragments per analyte and multiple isotopologs per fragment were considered. Thus, in order to account for the stable isotope variation, the correlation between fragment abundances was modified from a classical correlation among monoisotopic abundances to a correlation of the sum from all measured isotopologs for each fragment pair. Finally, for each amino acid analyte, three or more fragments were considered to provide a well-rounded annotation. From the final list of fragments, the most abundant ones (i.e., in the linear range of MS detection) were selected in order to calculate the percentage of ^15^N enrichment. Only fragments with null enrichment in the control were let to pass to the next stage. When all fragments presented residual “enrichment” in non-labelled samples this way considered a technical bias, and as such the mean “enrichment” in non-labelled samples was subtracted from the labelled samples and those fragments in which the final variance in control “enrichment” was minimal were further used. Finally, when multiple fragments satisfied the criteria to be useful as a proxy for amino acid synthesis, the most accurate were defined to be those fragments with the lowest relative standard deviation across technical triplicates and biological replicates (File S1). Subsequently, using the molecular formula information per fragment, NIA corrections and % of enrichment calculations (***Heinrich et al., 2018***) were performed, followed by a statistical comparison using the RandoDiStats R package (Tukey-HSD test was performed after an ANOVA with a confidence level of 95 %).

#### Plant Protein Synthesis Rates (K_*S*_)

Two main approaches were used to derive our own calculations of protein synthesis rates, which are both detailed below. In Ishihara et al., protein synthesis rates are calculated using the amino acid alanine and its enrichment percentage in proteinogenic (ALA_*p*_) and soluble (ALA_*S*_) pools as a proxy for label incorporation into protein (K_*S*_, equation 3a). The experimental time from the beginning of the experiment (t_0_) until the end (t_*f*_) is used to transform the nominal values into rates, which are expressed in percentage (%) per hour after a multiplication by 100.

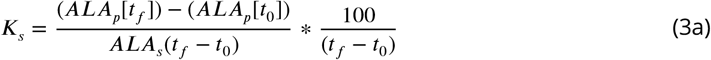

Protein degradation rates are then derived from K_*S*_ minus the product between plant relative growth rates (RGR) times the fractional change in protein content (P_*p*_), accounting for differential growth (K_*d*_, equation 3b).

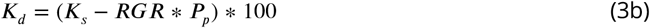

In Li et al., protein degradation rates are defined as the product of fold change in protein (FCP, equation 4c), natural logarithm normalized, by the non-labelled peptide fraction (1 - LPF) (K_*d*_, equation 4a). LPF corresponds to the integer of the area under the curve observed in isotopolog envelope shifts caused by tracer incorporation for specific peptides. The calculations are performed using an in-house script written in Mathematica and the percentages differ from the peptide enrichment values.

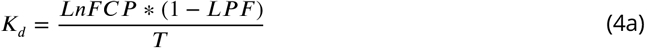

Protein synthesis rates are then derived as a product of FCP, weighed by the K_*d*_ and experimental time (T, 4d), times the K_*d*_ rate (K_*S*_, equation 4b).

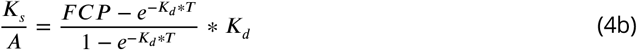

FCP is calculated as the ratio between the products of fresh weight (FW) times total final (P_*pf*_), in the numerator, and total initial (P_*p*0_) protein content in the denominator (FCP, equation 4c).

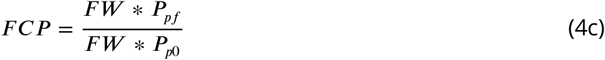

Time (T, equation 4d) equals to the difference between the intial experimental time (t_0_) and the final experimental time (t_*f*_).

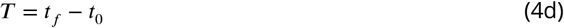

##### Measurement Units of Protein Synthesis Rates (K_*S*_)

Our K_*S*_ units imply percentage (%) of normalised labelled peptide fraction accumulated per unit of weight per hour (equation 5a).

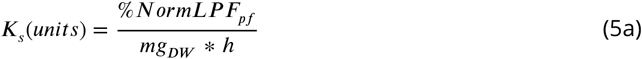

The product of the protein fraction (P_*f*_) times the labelled peptide fraction equals a normalised version of the labelled peptide fraction (NormLPF_*pf*_, equation 5b) that is turned into percentage fraction when multiplied by 100.

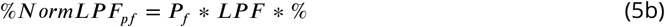

This operation intends to normalise the enrichment in single peptides by the fraction of accumulated total protein, preventing biases derived from differential protein accumulation. Ultimately, the NormLPF_*pf*_ units conserve the rate terms from the RGR formula, i.e., NormLPF_*pf*_ accumulated per unit of weight per hour (equations 5a & 5c).

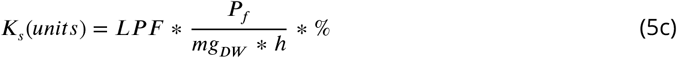

#### Ribosome Enriched Proteome

RAW chromatograms, including labelled and non-labelled samples, were processed with MaxQuant, version 1.6.10.43 (***Cox and Mann, 2008***). Search parameters included fixed—carbamidomethyl (C) and variable—oxidation (M), acetyl (protein N-term) modifications. Everything else was set as default. Subsequently all .RAW files were converted into .mzML using the MSConverterGUI from the ProteoWizard toolbox. The threshold peak filter was set to “absolute intensity”, ensuring the retention of all the peaks with an intensity greater than 100. This step allowed retaining in the files all the low abundant isotopes to conserve the isotopic envelopes of single peptides. Subsequently, using a python script, “isotopeEnrichment.py”, developed in-house (isotopeEnrichment), the abundances of individual isotopolog peaks for peptide signals were mined out of the .mzML files (Table S4 - D tab). Briefly, the sequence, mass, charge and retention times for peptides derived from proteins were taken from MaxQuant search results (i.e., the “evidence” MaxQuant output file). For each peptide localised at a single point in the chromatograms, theoretical exact masses were then used to create an extracted ion chromatogram (EIC) that is a summation of the intensities of each of the target isotopolog peaks. A Gaussian curve was then fitted to the EIC and the target isotopolog intensities were taken as the average of the observed intensities in a given number of mass spectra at either side of the Gaussian peak maxima. The procedure avoids skewing subsequent calculations to single scan isotopic ratios and thus ensures that the measured isotopolog abundances and relative ratios across the peptide peak are conserved with high fidelity. The script is written for python 3.7 and uses PymzML (***Bald et al., 2012***) and Pyteomics (***Levitsky et al., 2019***) to read mass spectrometry data files and calculate peptide masses respectively. The exemplary code used to produce our results was:

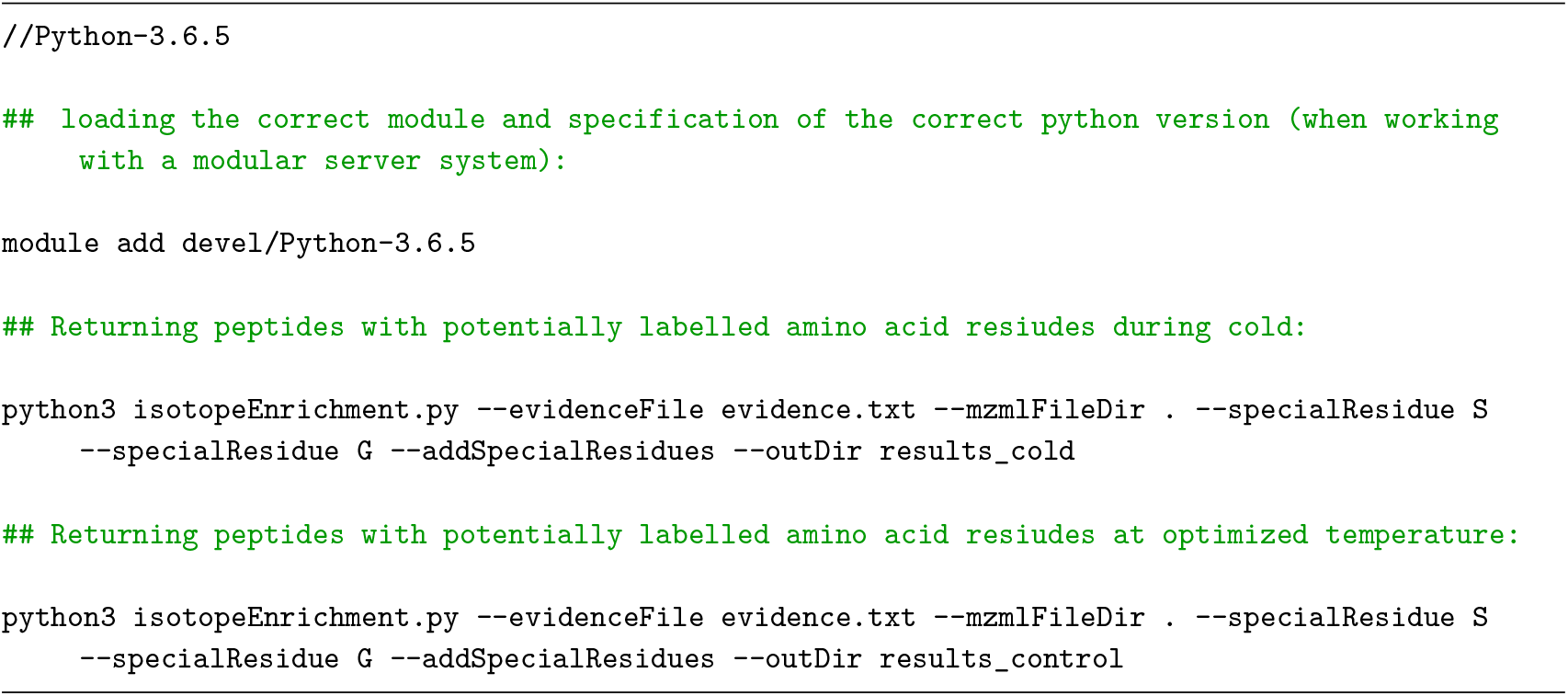

The mined isotopolog peaks for individual peptide species were then used to construct the necessary files for enrichment calculation (Table S4 - E tabs). Natural isotopic abundance (NIA) was calculated and abundances corrected using the R package IsoCorrectoR (***Heinrich et al., 2018***). The package requires first a “molecule” file that features the molecular formulas of the masses to be corrected. Secondly, an “intensities” file, containing the measured mass features and their isotopologs. Finally, an “elemental” file, containing the chemical elements with their NIA. Input files have been compiled in Table S4 - E tabs. An R function, “isotopeEnrichment.R”, was built to take up the abundances mined by “isotopeEnrichment.py” and turn them into the necessary format of IsoCorrectoR. Subsequently, the .csv files containing the enrichment percentages were used as input for the R function “EnrichmentSet.R”, which generates subsets of significantly labelled peptides using dependencies to the RandoDiStats R package. Additionally the function outputs the subset of unlabelled peptides as a control to monitor in the annotated proteins. The next function is named “AnnotateProteins.R”, which takes the outputs from “EnrichmentSet.R” in order to build protein enrichment percentages that are calculated based on the average enrichment of their monitored peptides. The function outputs protein sequences with the highlighted monitored peptides, the mean non-corrected protein enrichment percentage (non-corrected LPF, Table S4 - E4 tab) and standard deviations as a measure of reliability from the obtained averages. Finally, “LPFcorrect.R” corrects the outputs using the enrichment percentage in soluble amino acids and protein enrichments are transformed into fractional synthesis rates using equation 2a, all these calculations have been implemented into R functions and can be used from the ProtSynthesis R package. The workflow can be used averaging peptides from the same protein into a single entry or conserving peptides individually.

## Acknowledgments

We thank the Mass Spectrometry and Proteomics Facility of The Bio21 Molecular Science and Biotechnology Institute at The University of Melbourne for the support of mass spectrometry analysis. We thank Dr. Sneha Gupta for providing the seed material used in this work.

---

**Figure 1–Figure supplement 1.**
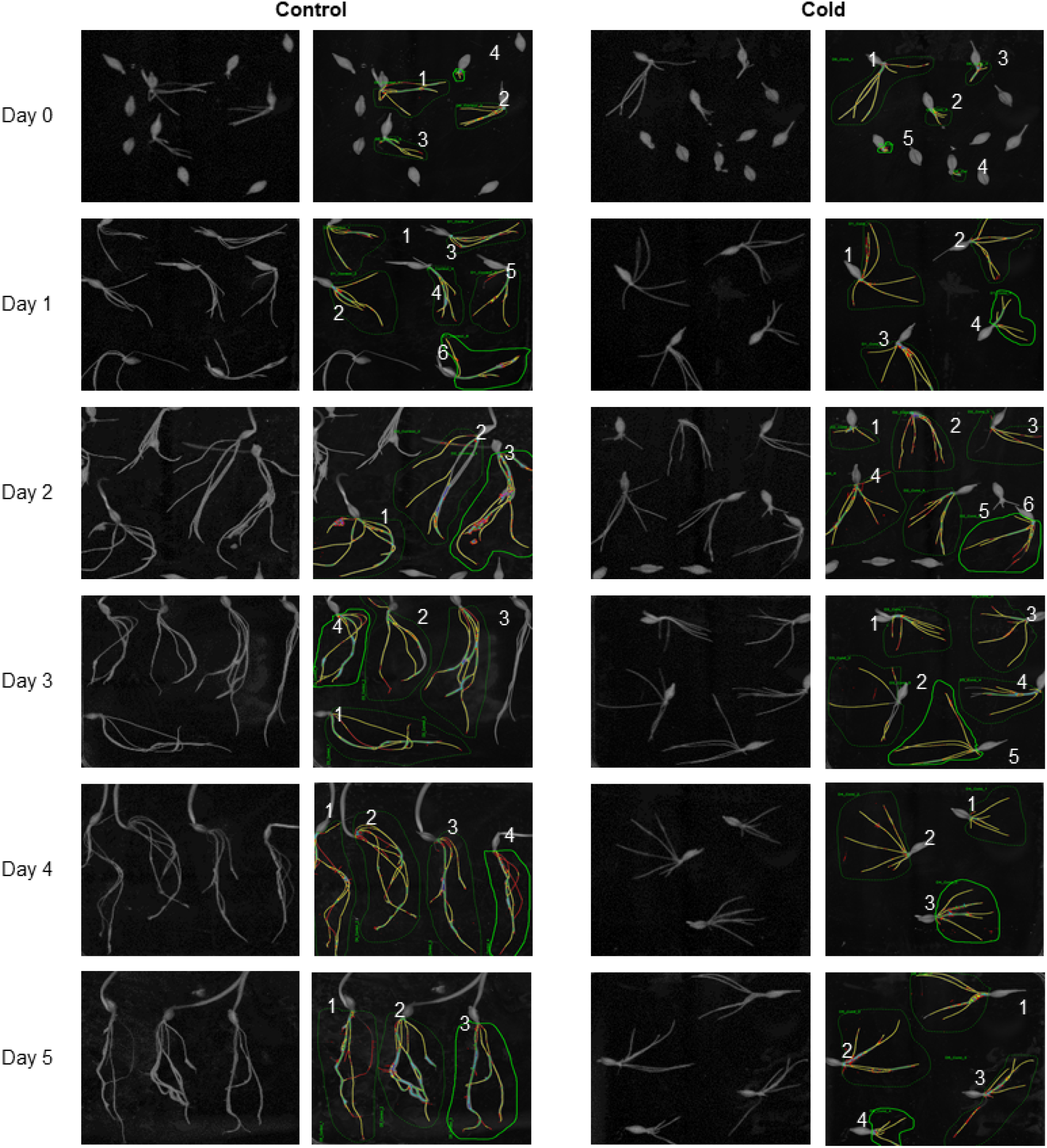
Root phenotype during germination assay in optimal control and suboptimal cold temperatures. Root systems were scanned at each time-point, beginning from day 0 until day 5. The columns on the left represent roots reared at optimal temperature and 3-6 selected representative root systems for phenotypic analyses The columns on the right represent the germination assay of roots acclimated to cold temperature during each time-point and 3-6 representative seedlings were phenotypically analyzed. winRHIZO software was used.

**Figure 1–Figure supplement 2.**
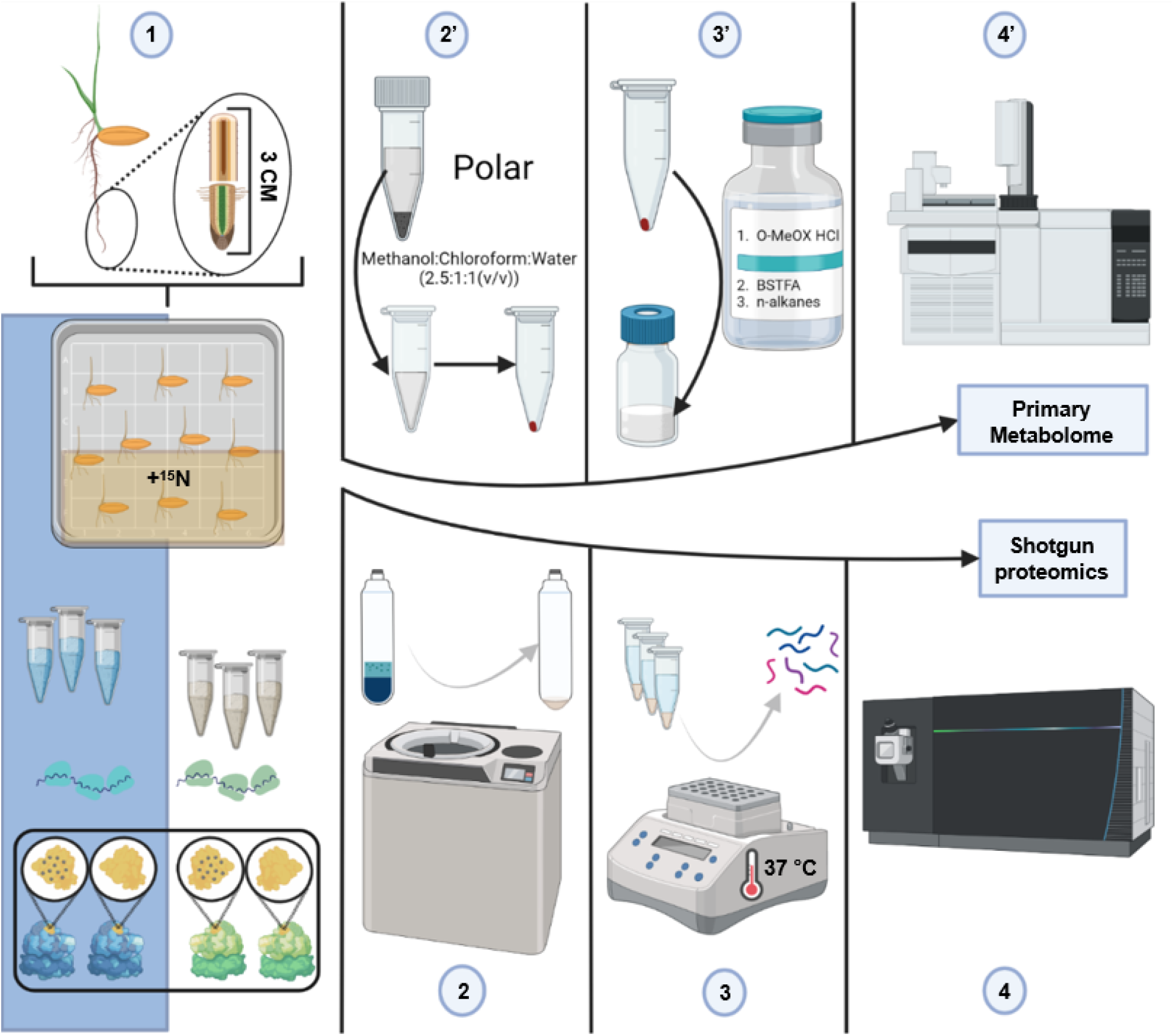
Summary of the methodological workflow to achieve measurements of protein synthesis and abundance in barley root tips. (1) harvesting of root tips from barley seedlings and division into two 1.5cm segments. Barley seedlings were germinated in two temperature regimes with one quarter of the plants having additional labelled nitrogen source and another quarter the same non-labelled nitrogen sources. (2) grinding of pooled tissue using liquid nitrogen, mortar and pestle followed by ribosome enrichment by ultracentrifuge-mediated large cellular complex subproteome extraction and pelleting. (3) Reduction, alkylation, trypsin digestion and peptide cleaning. (4) LC-MS/MS. (2’) Extraction of the polar primary metabolome using a methanol, chloroform, water ratio of 2.5:1:1 (V/V). (3’) Methoximation and silylation of primary metabolites. (4’) GC-ToF-MS multiplexed GC-APCI-MS.

**Figure 2–Figure supplement 1.**
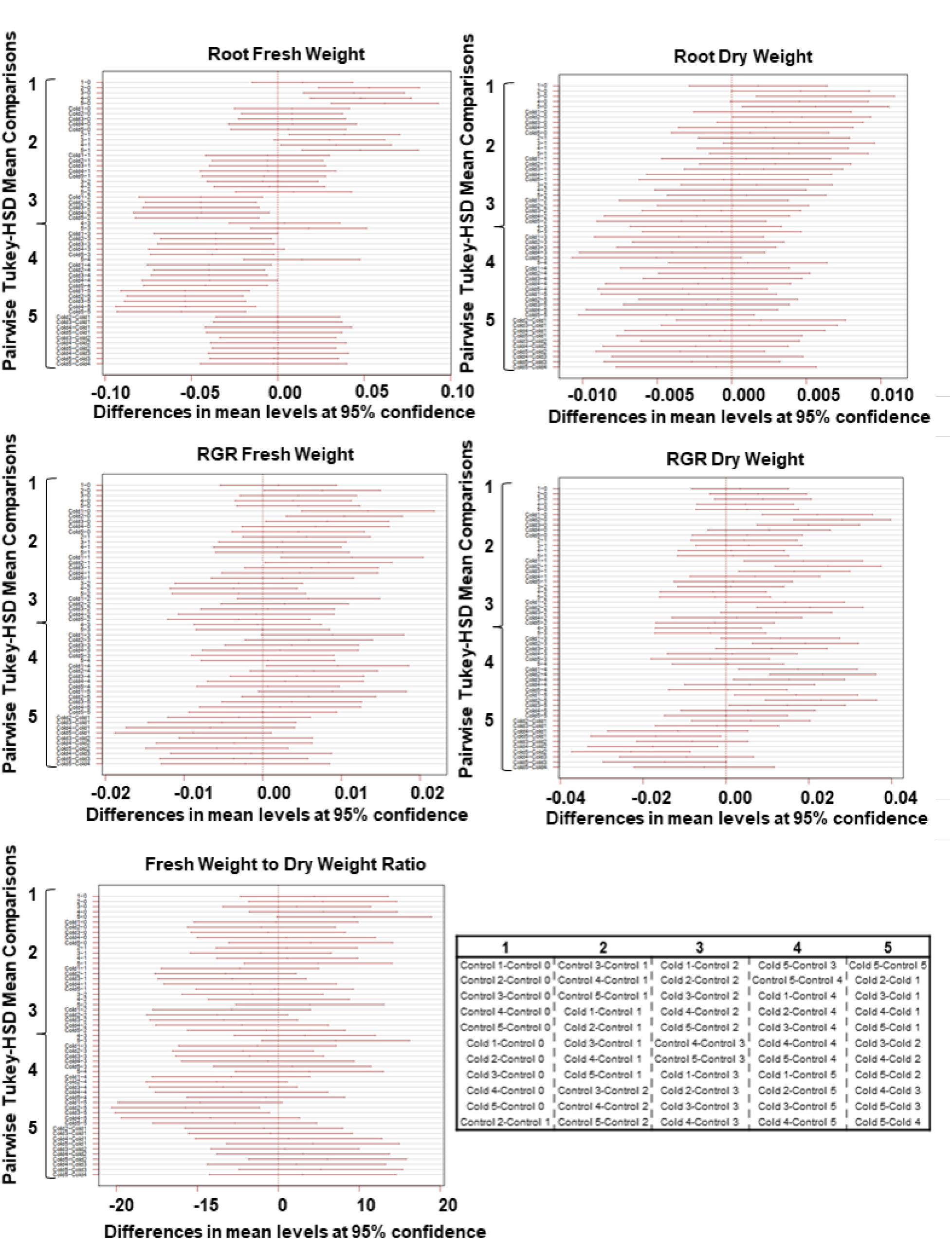
Summary of 95% confidence level Tukey-HSD statistical differences in mean levels of growth related variables across treated and control barley seedlings. Each panel reflects the pairwise comparison across all treatments of specific variables reflecting plant growth. The table on the lower right panel contains the sequential order of the mean comparisons in the plots in five groups. The groupings are signalled in the y-axes of the plots for reference.

**Figure 2–Figure supplement 2.**
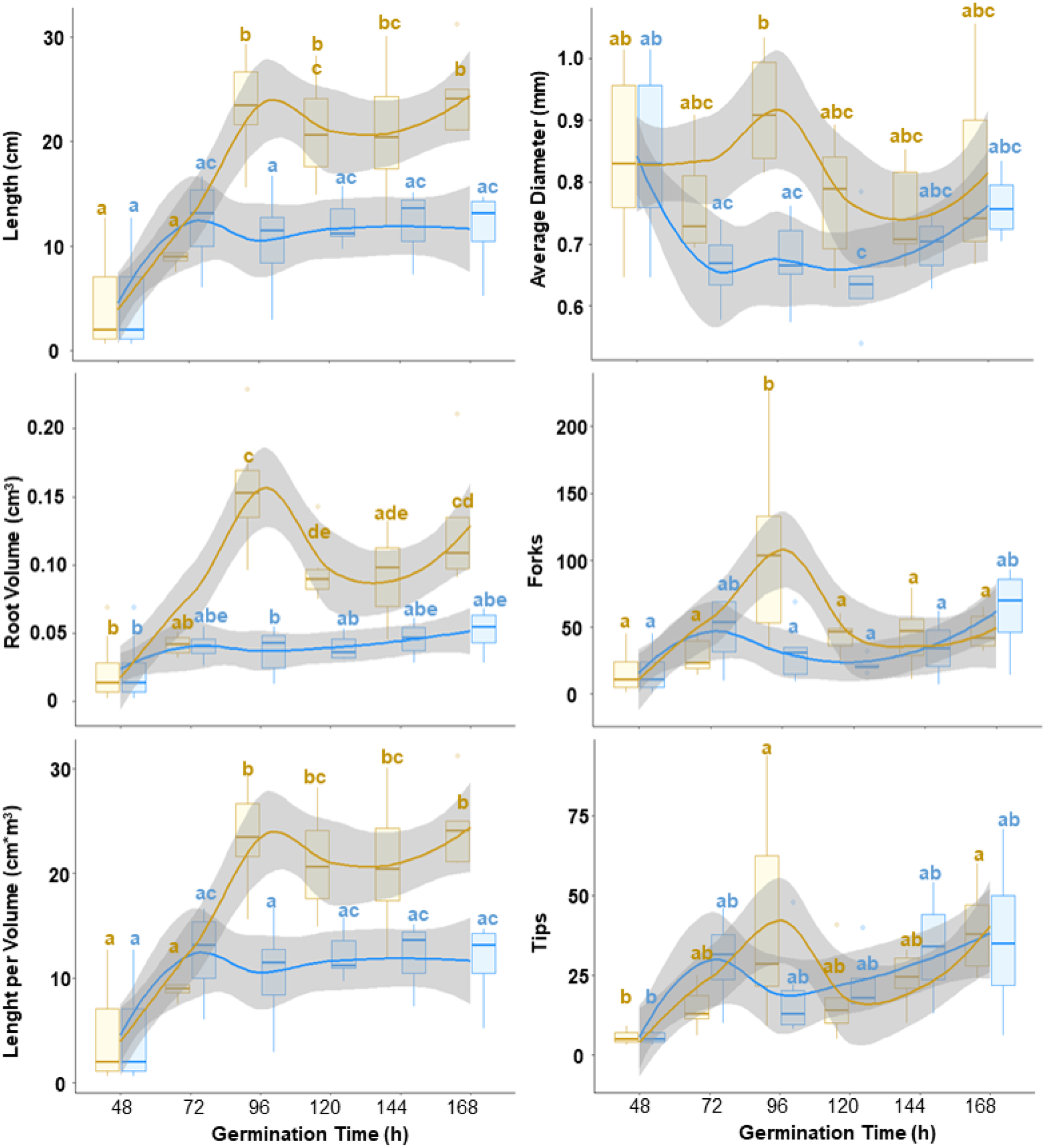
Statistical differences in mean levels (95% confidence level Tukey-HSD) of growth related variables derived from scanning treated and control barley seedlings at each time-point. Each panel reflects plant growth dynamics at control and suboptimal low temperature. All variables were measured using scanned images and the winRHIZO software.

**Figure 2–Figure supplement 3.**
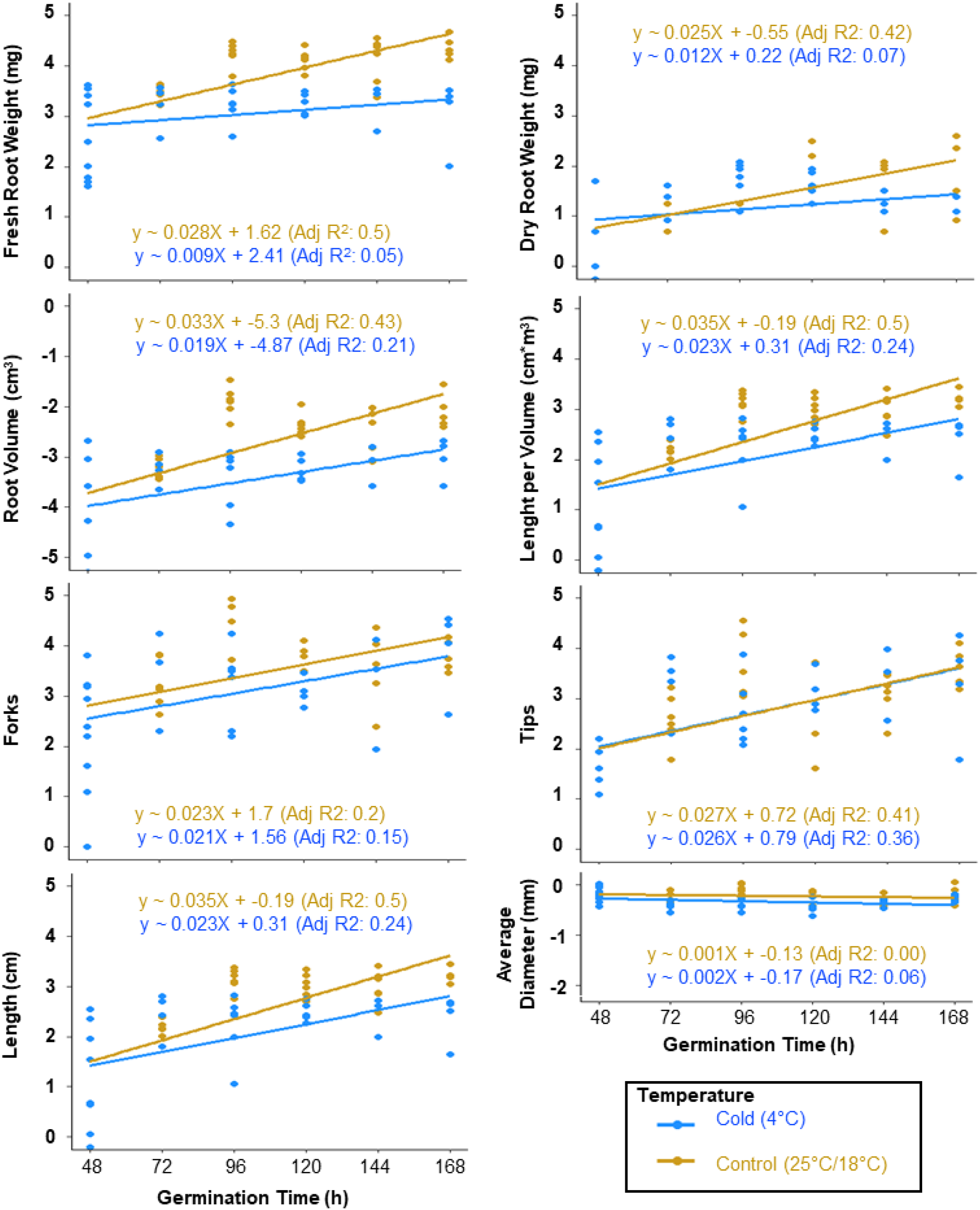
Linear regression after natural logarithm transformation of growth related variables. Growth variables were transformed using the natural logarithm (Ln) and a subsequent linear regression was made on the transformed vector. The fitting was evaluated using the adjusted r^2^, and the respective equation was derived from the linear fitting following the equation of a straight line (i.e., f(x) = mX + b, where m represents the slope and b the intercept). The slope represents mean growth rate for each variable and its biological accuracy depends on the adjusted r^2^ being close to 1.

**Figure 5–Figure supplement 1.**
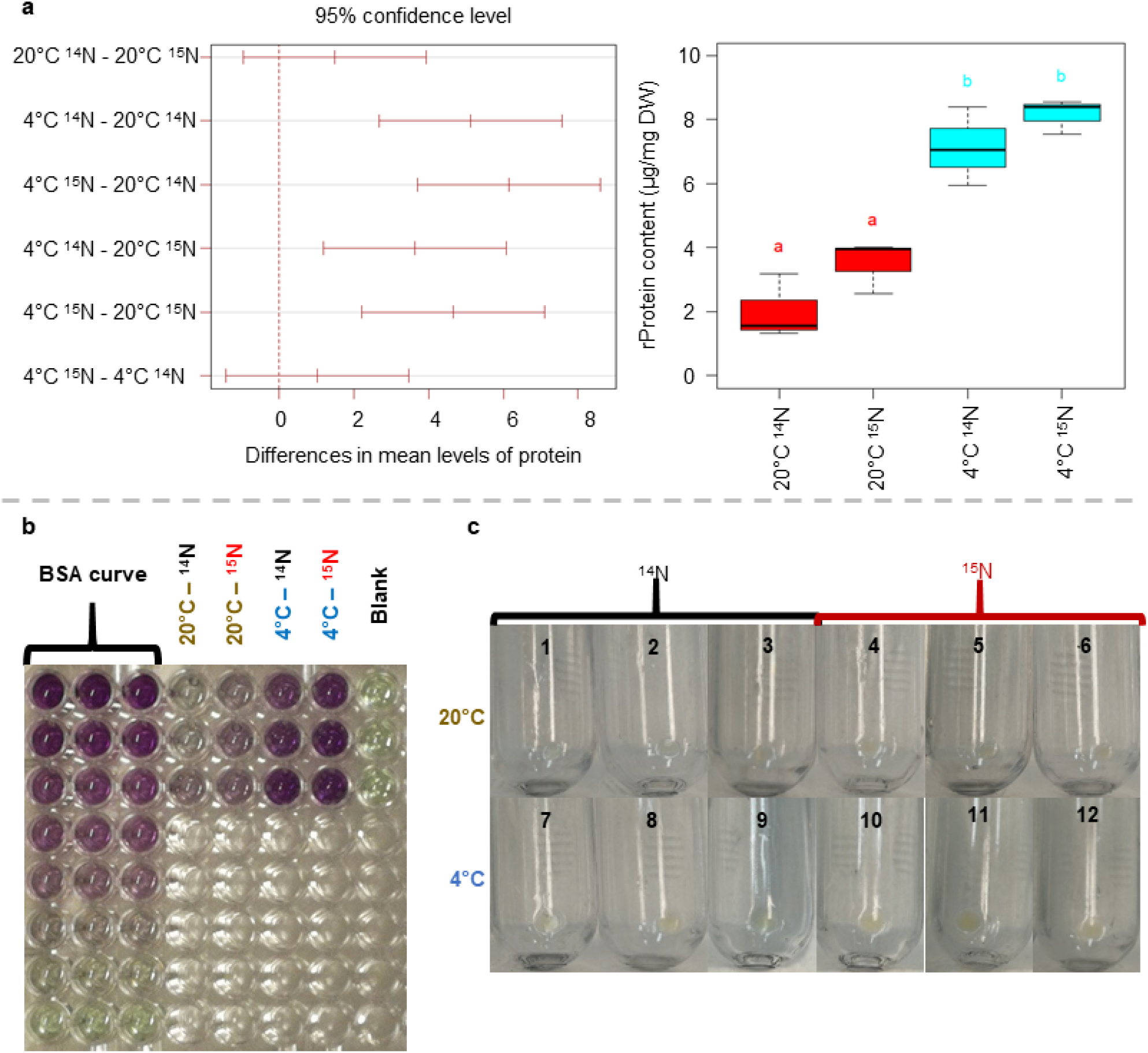
Summary of 95% confidence level Tukey-HSD statistical differences in mean levels of protein content from proteome fractions enriched in ribosomes across treated and control barley seedlings. Also relates to Table S5. (a left-panel) Pairwise comparisons across all treatments of a proteome fraction enriched in ribosomal protein content. The paired mean differences are signalled in the y-axis of the plot for reference. (a right-panel) Boxplot representation of the ribosomal protein content mean differences across temperature regimes, significance is signalled by colour transitions and different letters above the boxes. (b) Features the original photograph of the plate used for the bicinchoninic acid assay from which the results portrayed in panel “a” were derived. (c) Features the original photographs of the ribosome-enriched pellets from all experimental samples after passing through the 60 % sucrose cushion.

**Figure 5–Figure supplement 2.**
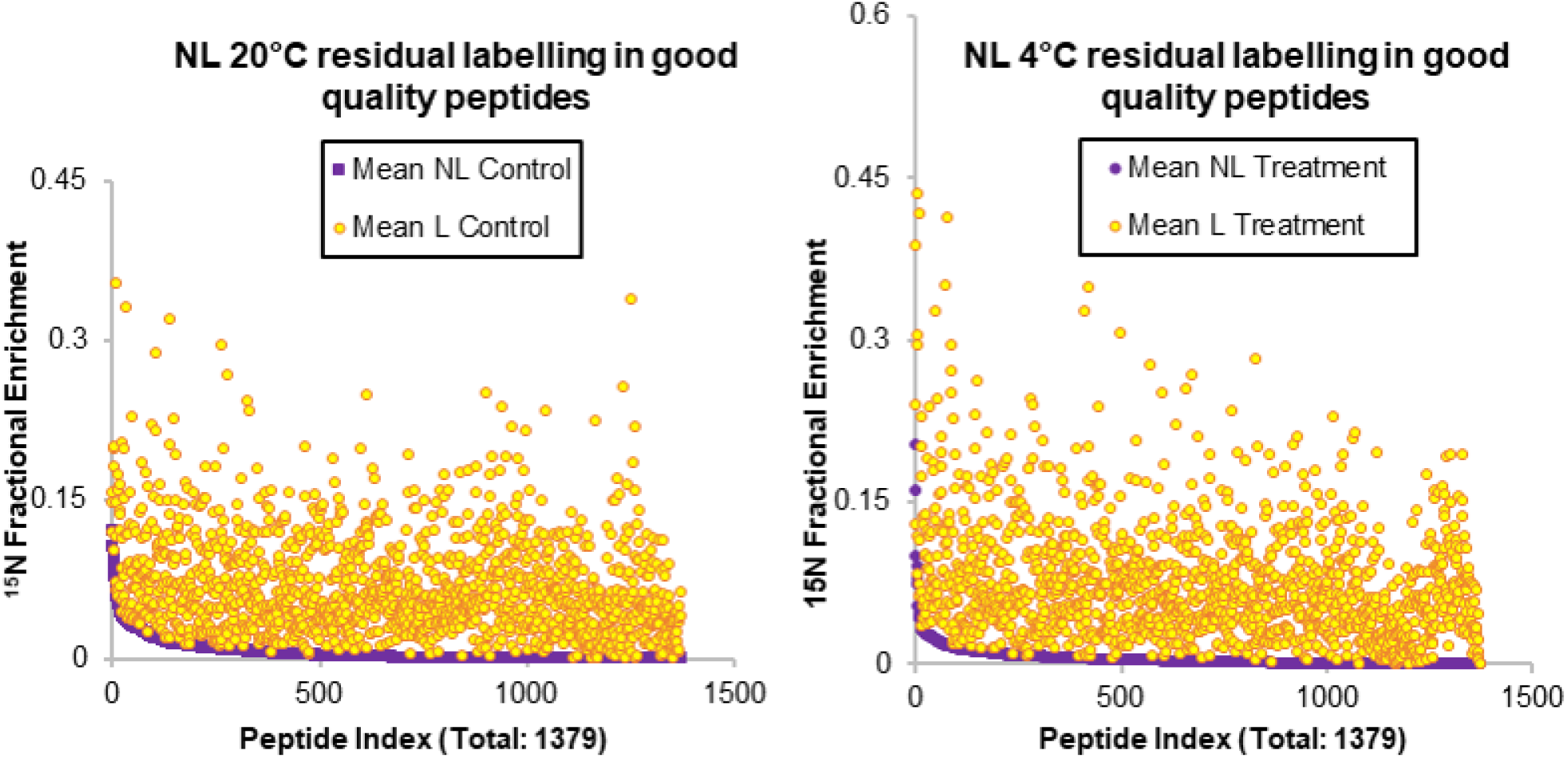
Subset of peptides considered as having optimal quality for interpretation of their relative fractional synthesis rates during the physiological transition of roots from germinating barley seedlings from optimal to sub-optimal temperature. The peptides in the list have all the necessary isotopolog abundances to calculate enrichment percentages in both temperature treatments. Many of the peptides in the list still conserve “noise” in the sense of false enrichment in the non-labelled samples (purple dots in both graphs). Nevertheless this noise is mostly below 1 % enrichment and always below the labelled samples. In both graphs the enrichment fraction is portrayed in the y-axis while the x.axis contains the peptide indexes, which have been sorted from highest to lowest noise in the non-labelled samples. Thus both graphs present a different peptide order. The left graph contains the information of peptides as monitored in samples derived from roots of germinating seedlings growing at an average of 22°C. The right graph contains the information of peptides as monitored in samples derived from roots of germinating seedlings growing at an average of 4°C.

**Figure 6–Figure supplement 1.**
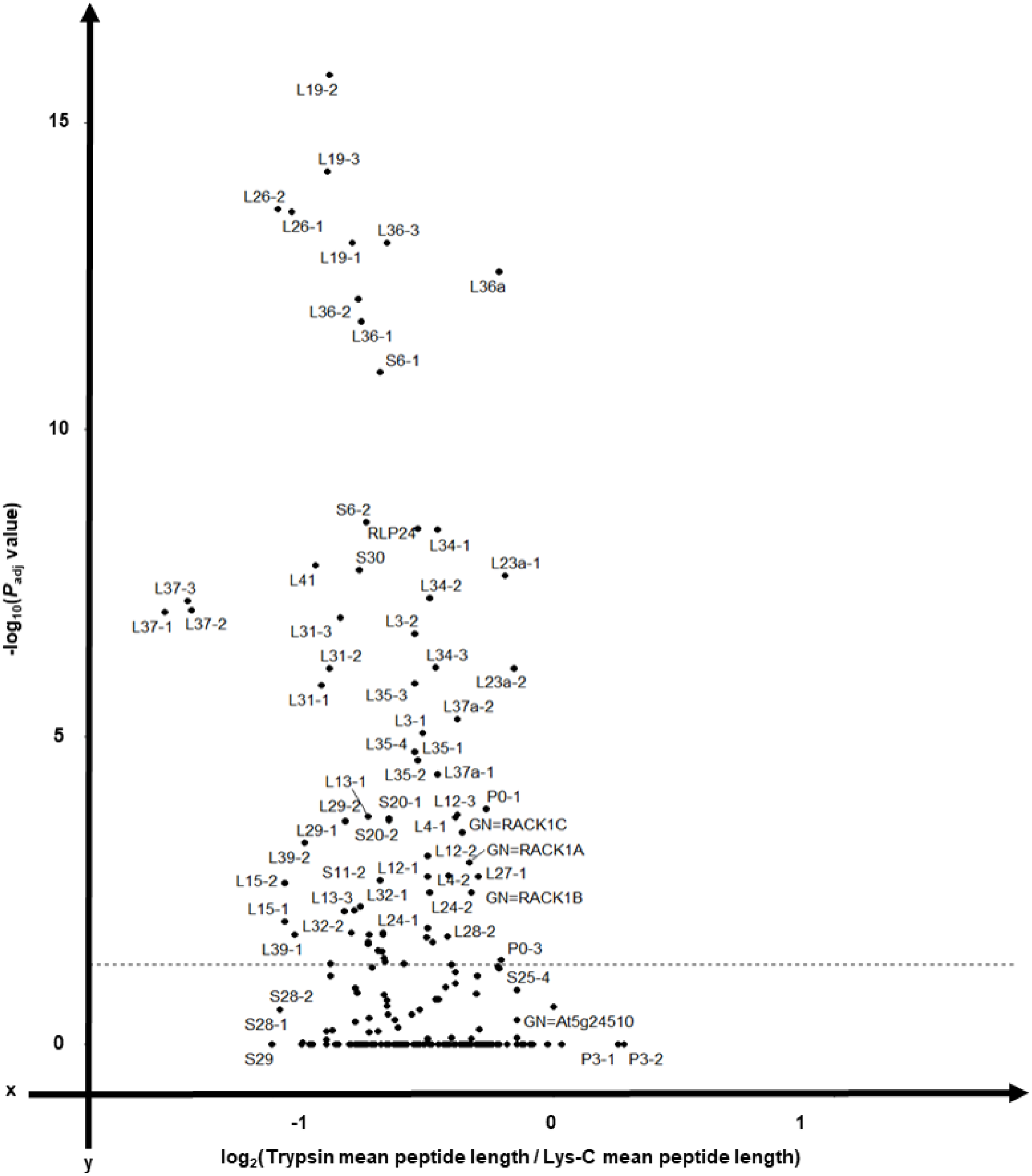
Volcano plot outlining the differences in mean peptide length produced with Lys-C or Trypsin, during an *in silico* protease digestion test, as proteases to digest the Arabidopsis signature plant ribosomal proteome. Also related to Table S6. Protease digestion *in silico* was performed with the free software Protein-Digestion-Simulator. The resulting plot from analysing the digestions contains in the x-axis the *log*_2_ of the ratio between mean peptide lengths from Trypsin and Lys-C digestion. Similarly, the y-axis contains the -*log*_10_ of the *P*_*adj*_ value from the statistical comparison of mean peptide lengths per each specific barley ribosomal protein. Note that with the exception of RPS21 and RPP3 all the other ribosomal proteins have shorter mean peptide lengths when digested with Trypsin as compared to Lys-C. The horizontal dotted line signals the significance boundary and thus all the proteins above this line have significantly shorter mean peptide lengths when digested with Trypsin as compared to Lys-C.

**Figure 6–Figure supplement 2.**
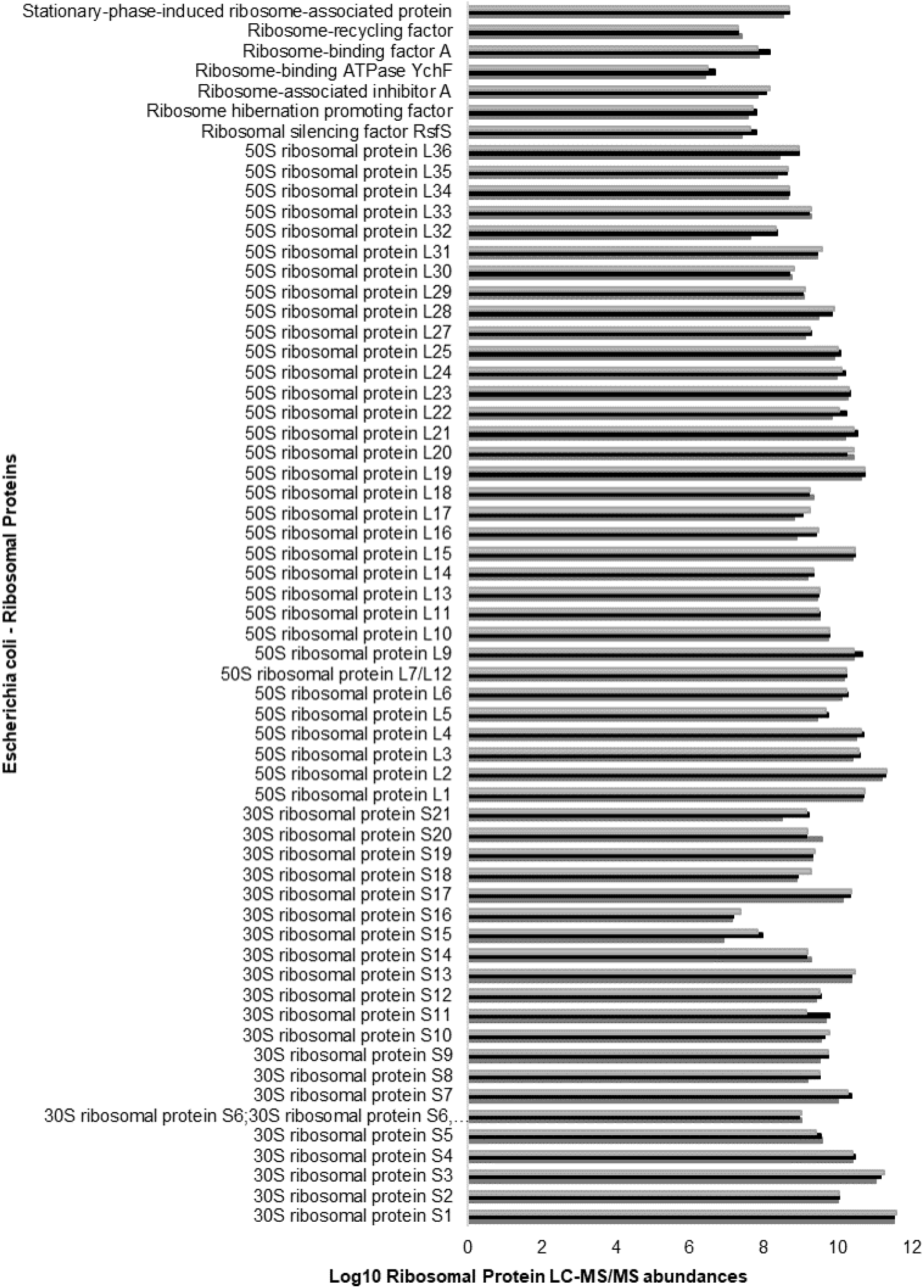
Positive control and complete coverage of *escherichia coli* 70S ribosomal proteome from a commercially available preparation used to verify the ribosomal proteomics pipeline. Also related to Table S7. The ribosomal proteomics pipeline was verified with three independent replicates from the same commercial preparation of *escherichia coli* 70S ribosomes (P0763S, NEB, Australia). The pipeline tested included ribosome extraction, subsequent purification through a sucrose cushion, resuspension of the pelleted complexes with a chaotrope to promote ribosomal protein dissociation and rRNA removal before SP3 beads binding of the ribosomal proteins for protease digestion. The coverage of the 70S ribosomes was full, with 21 proteins from the 30S small subunit and 33 from the 50S large subunit, plus a small set of ribosome associated factors. The height of the triplicate bars in the plot (i.e., x-axis unit) represents the *log*_10_ ribosomal protein abundances as measured by LC-MS/MS from the control ribosomal complexes. The y-axis contains the common name of the identified ribosomal proteins.

**Figure 6–Figure supplement 3.**
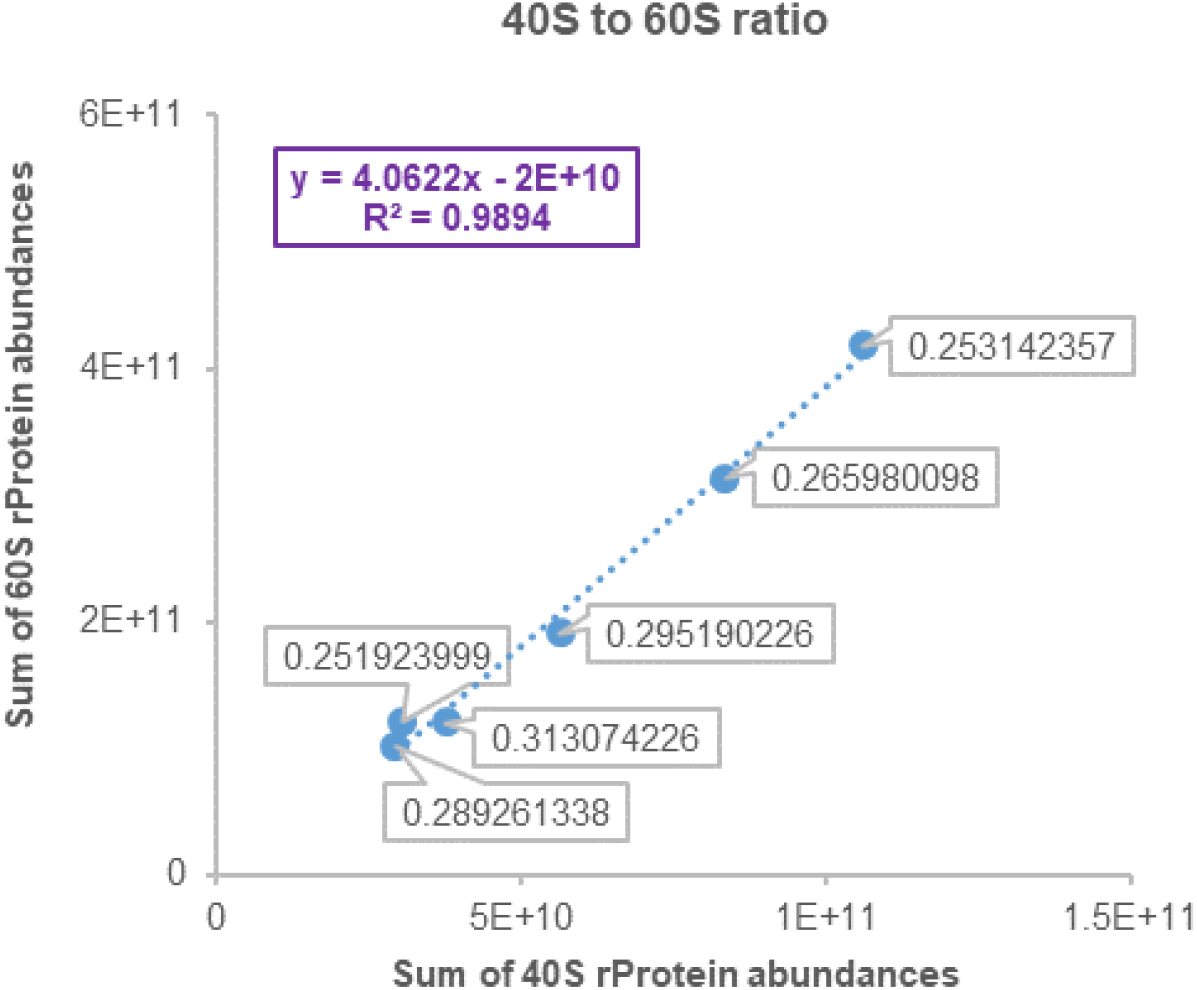
Ratio between 40S SSU and 60S LSU abundances across experimental samples. Also related to Table S4 - Tab C3. The y-axis features the sum of all detected and reliable 60S rProtein abundances. Likewise the x-axis features the sum of all detected and reliable 40S rProtein abundances. The labels at each point are the 40S to 60S ratio as calculated for each sample, note that the ratios all lie between 0.25 and 0.31, which means that 60S proteins are always at a relationship of 3 to 1 with 40S proteins and this aligns with the number of protein paralogs per subunit. Finally, since the ratios are constant, note that the dispersion of the dots is well adjusted to a linear regression model with an r^2^ above 0.98.

